# A bipolar taxonomy of adult human brain sulcal morphology related to timing of fetal sulcation and trans-sulcal gene expression gradients

**DOI:** 10.1101/2023.12.19.572454

**Authors:** William E. Snyder, Petra E. Vértes, Vanessa Kyriakopoulou, Konrad Wagstyl, Logan Z. J. Williams, Dustin Moraczewski, Adam G. Thomas, Vyacheslav R. Karolis, Jakob Seidlitz, Denis Rivière, Emma C. Robinson, Jean-Francois Mangin, Armin Raznahan, Edward T. Bullmore

## Abstract

We developed a computational pipeline (now provided as a resource) for measuring morphological similarity between cortical surface sulci to construct a sulcal phenotype network (SPN) from each magnetic resonance imaging (MRI) scan in an adult cohort (N=34,725; 45-82 years). Networks estimated from pairwise similarities of 40 sulci on 5 morphological metrics comprised two clusters of sulci, represented also by the bipolar distribution of sulci on a linear-to-complex dimension. Linear sulci were more heritable and typically located in unimodal cortex; complex sulci were less heritable and typically located in heteromodal cortex. Aligning these results with an independent fetal brain MRI cohort (N=228; 21-36 gestational weeks), we found that linear sulci formed earlier, and the earliest and latest-forming sulci had the least between-adult variation. Using high-resolution maps of cortical gene expression, we found that linear sulcation is mechanistically underpinned by trans-sulcal gene expression gradients enriched for developmental processes.

## Introduction

The adult human brain has a complex, wrinkled surface formed by undulation between grooves (sulci) and ridges (gyri) of the cortical sheet. The cerebral cortex forms embryonically from distension of the neural tube, and its surface remains smooth (lissencephalic) until about 20 weeks gestational age (GA) in humans^1,2^. In the second half of pregnancy, the cortical sheet becomes progressively more wrinkled (gyrencephalic), as a growing number of sulci appear to indent its surface^3–5^. At birth, the sulco-gyral patterning of the brain is thought to be close to its life-long final configuration^6–8^, as complexly patterned and individually unique as the fingerprints that are also enduringly formed at birth^9^. However, it has been challenging to quantify such a spatio-temporally complex and individually variable process as human brain cortical surface patterning.

Existing sulcal taxonomies are generally descriptive and anatomically referenced to surrounding cortical areas (*e.g.*, central sulcus or parieto-occipital fissure) rather than the size or shape of the sulci themselves. Expert but largely qualitative postmortem examinations of limited quantities of human brains have supported the classification of adult sulci into two or three categories or classes, linked to their timing of fetal emergence or sulcation^3,4^. These classifications - into primary, secondary, or tertiary sulci - have been defined differently across studies but can converge on notions of time of sulcal emergence, hierarchical relationships in sulcal size, and phylogenetic presentation^3,10,11^. Sulci consistently classified as primary, like the central sulcus, are the first and longest sulci to form and are conserved in shape across primates^12^. In humans, primary sulci begin to appear in the second trimester of pregnancy, from about 20 weeks GA, until

∼32 weeks GA (early third trimester). Secondary sulci are sometimes defined as the more variable branches from primary sulci^3,8^ and emerge between ∼32 and 38 weeks GA. Alternatively, these sulci are defined as separate sulcal regions forming in this latter developmental period^13^. The few tertiary sulci similarly emerge after 38 weeks and for some months post-natally^2,3^. Annotations of adult cortex have offered an overlapping classification scheme^14^ which defines sulci by their length and continuity, providing an atlas of possible configurations of secondary and tertiary sulci. However, this primary-to-tertiary labeling model remains quite arbitrary with considerable uncertainty over the correct labeling for many individual sulci (*e.g.*, the middle frontal sulcus is assigned a different label for each of Chi et al.^3^, Dubois et al.^8^, and Voorhies et al.^13^). To date, there has been a dearth of systematic quantitative frameworks for estimating relatedness between different folds as a foundation for sulcal classification.

Complementary to taxonomic studies, biomechanical models have focused on predicting what regions of cortex become sulcus versus gyrus during development as opposed to their prospective primary versus tertiary sulcal patterning^15–17^. These models have identified genes whose mutations are thought to play a causal role in cortical malformation disorders^18,19^. But, these models and biophysical simulation studies^20,21^ presently remain incomplete, unable to explain sources of individual variability in sulcal patterning or the protracted regional differences in the timing of sulcal formation^5^. Fundamentally, our understanding of sulcal complexity and its ontogenesis in fetal life has been limited by the scale and quantitative scope of postmortem brain studies, which lack sufficient sample size to encompass high levels of between-subject variability in later-forming sulci^22,23^.

The advent of computational methods for sulcal morphometry from magnetic resonance imaging (MRI) data, *e.g.*, BrainVISA’s Morphologist toolbox (http://brainvisa.info)^24–26^, has advanced the field by standardizing and automating sulcal segmentation, enabling fully quantitative assessments of sulcal shape. Geometric measurements, such as surface area and depth, along the length or fundus of a sulcus, are heritable^23,27^, related to cognition^13,28^, and related to psychiatric and neurodevelopmental disorders^27,29^. Measures driven by sulcal patterning such as global and local gyrification^30–32^, as well as cortical surface area and curvature^33,34^, have also pointed to the importance of sulcal patterning in health and disorder. Contemporary methods of computational sulcal morphometry have enabled automated, reliable, whole brain quantification of sulcal size and shape^26,35^, allowing more detailed assessments of individual differences and genetic effects on adult morphological phenotypes measured at each of multiple sulci^23,27^. Although there is increasing interest in measuring the similarity of cortical areas in terms of multiple MRI measures of geometry and tissue composition, using methods like morphometric similarity network (MSN) analysis^36^, there have been very limited comparable investigations of systemic patterns of covariation between sulci in terms of their size and shape metrics. Additionally, recent advances in analysis of complex sulcal anatomy from adult MRI scans have not yet been linked to insights into the developmental timing of sulcation, as measured in fetal MRI data.

In this context, we integrate two complementary technical innovations to define a new taxonomy of human cortical sulcation: (i) sulcal phenotype network analysis and (ii) growth curve modeling of fetal sulcation. We show that this new taxonomy of cortical sulcation in adulthood can be linked to the timing of brain sulcation in development and is coupled with diverse facets of multiscale cortical organization spanning from gene expression gradients to functional brain networks.

First, we developed sulcal phenotype network analysis from five key developmentally salient sulcal phenotypes (median and variability of sulcal depth, longest branch length, branch span, and fractal dimension) measured for each of 40 sulci across both cerebral hemispheres in a large sample of adult brain MRI scans (N∼35,000; UK Biobank)^37,38^. Pairwise sulcal similarity defined scan-level sulcal phenotype networks, which could be dimensionally reduced by principal component analysis to a bipolar axis ranging from linear (at the negative pole, -1) to complex or fractal sulcal geometry (at the positive pole, +1). To directly assess the origin and development of sulcal similarity, we aligned annotated maps of adult sulcal anatomy to a large fetal sample (N=228) of cortical surface reconstructions produced by the developing Human Connectome Project (dHCP)^39^. We used the well-studied mean curvature metric^40^ to map fetal sulcation trajectories for each of 40 adult brain sulci to promote a standard methodology amongst the often qualitative and sometimes inconsistent estimates of when each sulcus forms^2,5^. Strong coupling was observed between adult morphology and fetal development, with linear sulci forming before complex sulci *in utero* and the variability of linear or complex morphology being reduced at both the earliest and latest stages of sulcal development. Finally, we leveraged transcriptomic data to examine the mechanistic hypothesis that the most heritable, earliest-forming, linear sulci arise along boundaries of sharp transitions in cortical composition. We used dense expression maps of 20,781 genes^41–43^ to identify trans-sulcal gradients of gene transcription that are aligned orthogonally to the fundus of highly linear sulci in adulthood, and show that these genes are enriched for neurodevelopmental and developmental disorder gene ontologies. Together, our resource comprises comprehensive structural, functional, adult maturational, fetal developmental, and transcriptional annotation of sulci to enable future works to recognize and integrate our new taxonomy in fetal and adult human neuroscience.

## Results

### Sample

Adult brain structural magnetic resonance imaging (sMRI) data were obtained from the second release of the UK Biobank sMRI cohort^37,38^. A single sMRI scan was available for each of N∼39,000 participants (mean age = 64 years; 47% male) and was processed with FreeSurfer (http://surfer.nmr.mgh.harvard.edu/)^44,45^ and BrainVISA Morphologist (http://brainvisa.info)^24–26^ software. After excluding participants with neurological diagnoses^46^ (N = 1,568) and scans that did not pass quality control (N = 2,247, see **Methods**), we retained an analyzable sample of N = 34,725 scans (mean age = 64 years; age range = 45-82 years; 46% male) (**Figure S1**).

Fetal brain sMRI data were collected and processed with the developing Human Connectome Project (https://www.developingconnectome.org/) structural pipeline^39,47^. A total of N = 228 fetuses aged 21-36 gestational weeks were scanned *in utero* (**Figure S1**) and passed quality control for reconstruction of the inner cortical surfaces bilaterally (**Methods**).

### Sulcal phenotypes

For each adult brain scan, we segmented and labeled 40 sulci (20 per cerebral hemisphere [**Figure 1A**], see nomenclature in **Figure S2**) and estimated 5 shape metrics or features for each sulcus (**Figure 1B**): average (median) and variability (median absolute deviation) of sulcal depth; longest branch length; branch span; and fractal dimension (FD). These five measures were chosen to encompass morphometric variation both tangential and radial to the cortical surface. Prior studies have examined these features in isolation from each other^3,13,14,28,48–52^ but have not harnessed them as a combined group that can collectively triangulate the complex topography of any given sulcus.

**Figure 1.**
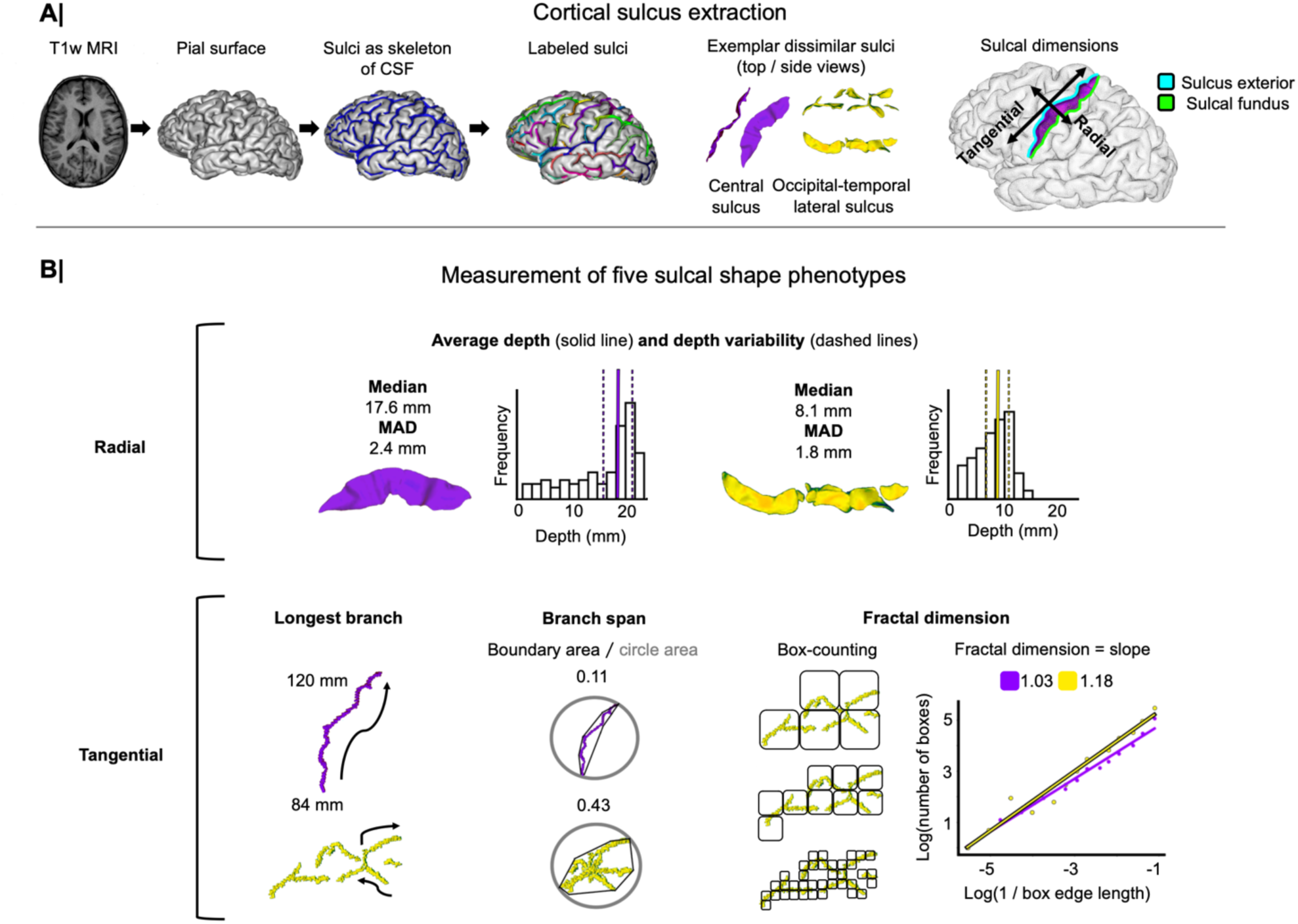
A suite of sulcal morphometrics spanning radial and tangential dimensions. (**A**) Using FreeSurfer and BrainVISA software, 20 sulci per brain hemisphere were segmented and labeled in each of 34,725 adult brain MRI scans. Sulci could then be measured along the dimension radial (from sulcal fundus to sulcus exterior) or tangential to the cortex (along the sulcus exterior). (**B**) Five shape metrics were measured for each sulcus, spanning the radial and tangential directions. Two metrics measured radial dimensions of sulci: average or median depth of the sulcal floor and variability of the depth of the floor along the length of the sulcus. Three metrics measured tangential dimensions of sulci: the length of the longest branch, branch span and fractal dimension (FD). Depth metrics were based on depth histograms, computed as the distance between points along the bottom of a sulcal valley (sulcal fundus) to the enclosing surface encompassing gyral peaks. The median was estimated as a measure of central location and the median absolute deviation (MAD) as a measure of variability. The length of the longest branch was the maximum geodesic distance spanned by any branch of a sulcus. Fractal metrics captured the space occupancy of sulcal shapes tangential to the cortex. Branch span was estimated by drawing a convex hull around each sulcus projected into 2D, and calculating the ratio of the area enclosed by the hull to the area of a circle that circumscribes the hull, 0 < branch span < 1. More complex, densely branched sulci occupy more of the space and have higher branch span than more linear, sparsely branched sulci. FD was estimated by a box-counting algorithm: the sulcus was iteratively tiled with boxes of different sizes and the number of boxes occupied by the sulcus at each size was counted; then ∼1 < FD < 2 was estimated by gradient of the log-log linear relationship between the size of boxes and the number of occupied boxes. More linear sulci have FD close to the limit of a Euclidean line (1) and more branched or complex sulci have FD closer to the limit of a Euclidean plane (2).

Plots of average sulcal phenotypes had similar cortical patterning across phenotypes (**Figure 2A**), indicating coordination between phenotypes within subjects. The {1 x 40} feature vectors describing each phenotype’s distribution across 40 sulci within an individual brain were correlated between the five sulcal phenotypes and averaged across subjects. Average sulcal depth and depth variability were strongly positively correlated with each other, and with longest branch length; whereas branch span was negatively correlated with sulcal depth metrics and positively correlated with FD (**Figure 2B**, left). For example, the central sulcus and parieto-occipital fissure were both deep and linear in terms of low branch span and fractal dimension. These within-brain correlations were consistently expressed across subjects (**Figure 2B**, right).

**Figure 2.**
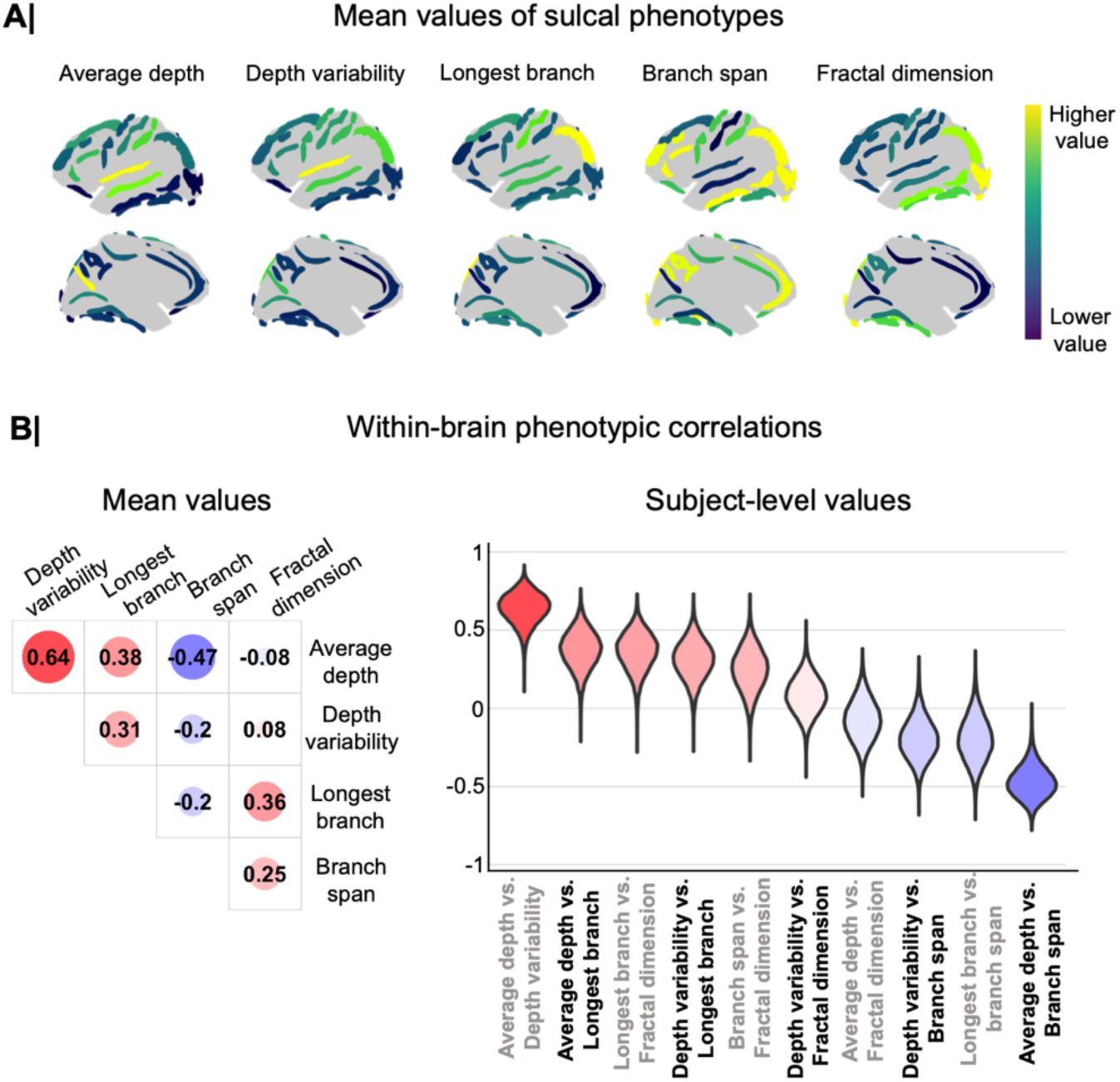
Cortical surface patterning of sulcal phenotypes and relationships between phenotypes. (**A)** Mean values of sulcal phenotypes are plotted on a scale highlighting relative differences between sulcal regions. The colors are shown on average sulci models, delineating their typical path between gyri. The gradients of sulcal anatomy are largely similar across the phenotypes, with sulci at the extreme ends of one phenotype often occupying extreme ends of other phenotypes. (**B)** Average correlation between {1 x 40} sulcal phenotype vectors within brains in the adult UK Biobank dataset (left) and distribution of these correlations across individuals (right). Sulcal phenotypes exhibit a variety of inter-measure correlations, highlighting the shared and complementary information afforded by each feature.

Alternatively, sulcal covariation between subjects was measured as the correlation between the pair of {1 x 34,725} feature vectors representing a single phenotype, *e.g.*, sulcal depth, at each of two sulci, *e.g.*, central sulcus and lateral fissure average depth. In contrast to the strong covariation across folds within subjects, there was low covariation between folds across subjects (median absolute correlation = 0.03) (**Figure S3**), in line with findings from Sun et al. (2022). Between-subject covariation was further reduced when first correcting sulcal phenotypes for linear effects of age, sex, and total brain volume (TBV) (correlation of residualised phenotypes = 0.01) (**Figure S3**). In exception, sulcal covariation across subjects was low but relatively higher between phenotypes for the same sulcus (block-wise diagonals in **Figure S3**) (median absolute correlation = 0.17, and after residualization = 0.16). Thus, it appears that different sulci show broadly reproducible morphological motifs across individuals (*e.g.*, deep and linear vs. shallow and complex), but substantial individual differences in sulcal morphology result in generally low between-subject covariation.

We investigated the effects of total brain volume (TBV), sex, and age on each phenotype at each sulcus (**Figure S4A**). Covariates had significant effects on multiple different sulci but overall had small effect sizes (mean absolute standardized effect size: age = 0.022, sex = 0.080, TBV = 0.163). We report this extensive covariate profiling of sulcal phenotypes in **Table S1**, providing standardized beta coefficients and significance for each of 40 sulci, 5 sulcal phenotypes, and 3 covariates. Using these covariate residualized phenotypes to repeat the analyses shown in **Figure 2B** revealed that within-brain sulcal patterning relationships were almost identical after removing age, sex, and TBV effects (**Figure S4B**).

### Sulcal phenotype networks (SPNs) reveal two dominant modes of sulcal morphology

To further investigate the similarity or dissimilarity between each pair of sulci in each adult brain, we estimated the {40 ⨉ 40} matrix of Pearson’s correlations between the feature vectors of the same 5 metrics measured at each sulcus, for each brain scan. This matrix was designated the sulcal phenotype network (SPN), with individual SPNs averaged to estimate the group mean SPN (**Figure 3A; Table S2**).

**Figure 3.**
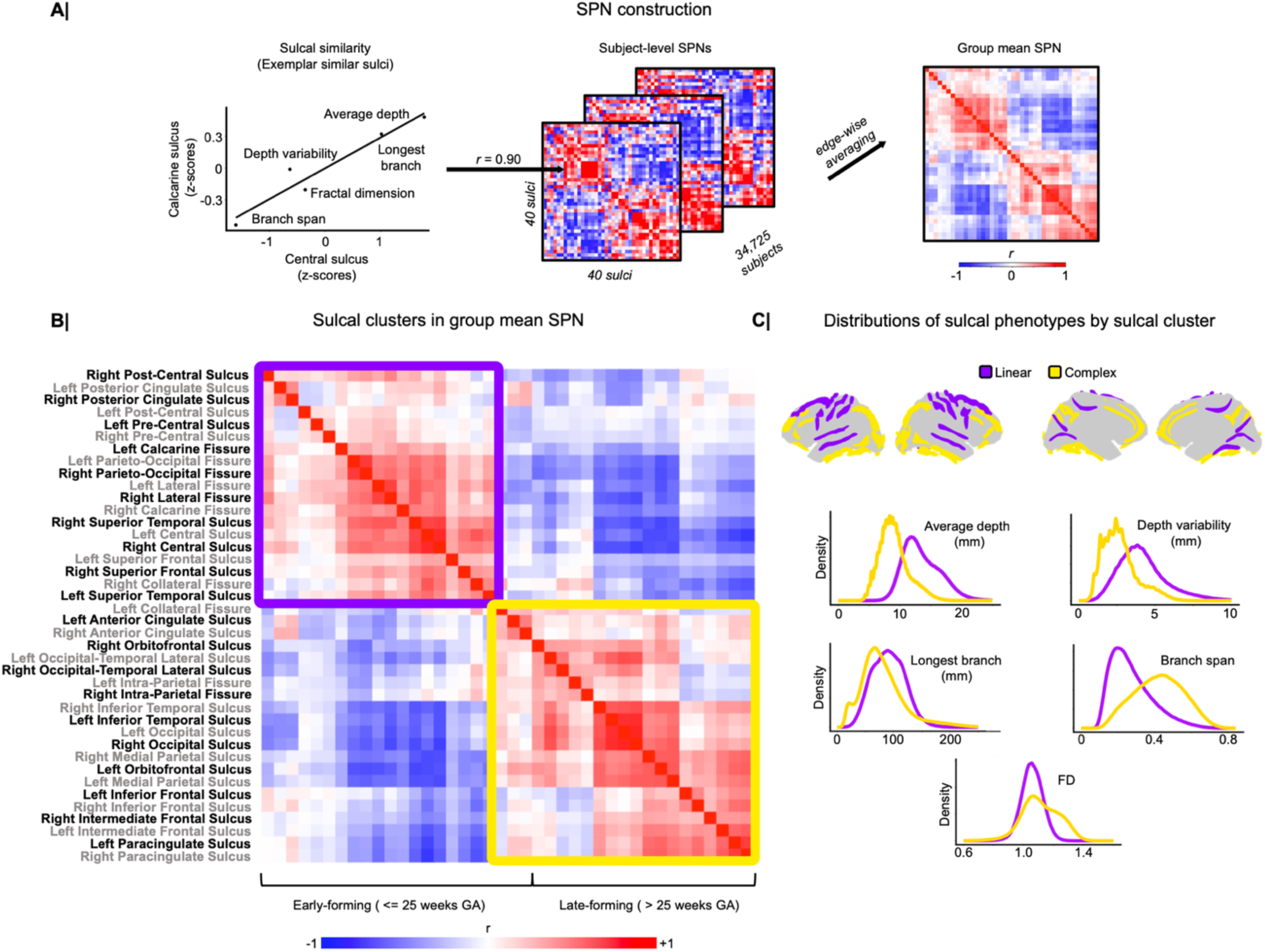
Sulcal phenotype network (SPN) analysis demonstrates that adult sulcal patterning is categorically represented by two major clusters, each comprising sulci which share distinctive (linear or complex) structural phenotypes. (**A**) Pearson’s correlation between the five sulcal phenotypes measured at each of a pair of sulci (Z-scored within each brain per metric) was used as an estimator of the morphometric similarity of two sulci within the same brain. Repeated for each possible pair of 40 sulci, this analysis generated a {40 ⨉ 40} correlation matrix which represents the similarity (r > 0) or dissimilarity (*r* < 0) of sulcal phenotypes across the whole brain, and this was designated the sulcal phenotype network (SPN). Edge-wise averaging across subjects yielded the group mean SPN. (**B**) The group mean SPN had two large clusters, separated almost entirely by early (<=25 weeks GA) and late (>25 weeks GA) forming sulci. (**C**) The cluster termed “linear” was generally located in more central regions on the lateral and medial faces of the brain with “complex” sulci situated at more extreme anterior, posterior, or ventral locations. Linear and complex names for sulcal clusters were based on the distinctive distribution of sulcal phenotypes displayed by each cluster, with the linear sulci having higher values in linear measures and complex sulci having higher values in fractal measures.

Hierarchical cluster analysis of the group mean SPN highlighted two large clusters of sulci that were positively correlated with (or similar to) other sulci in the same cluster; and negatively correlated with (or dissimilar to) sulci in the other cluster (**Figure 3B**). The two clusters were also clearly differentiated in terms of the characteristic morphometry of their constituent sulci: one cluster, designated *linear*, comprised sulci with greater mean depth, variability of depth, and longest branch length; the other cluster, designated *complex*, comprised sulci with greater branch span and fractal dimension (**Figure 3C**). Based on previous, postmortem labeling of when fetal sulci emerge^3^, we identified that the linear cluster’s sulci formed almost exclusively by or before 25 weeks GA and the complex cluster’s sulci formed almost exclusively after 25 weeks GA (**Figure 3B**). The clustering was further validated by the Dunn Index, finding hierarchical clustering was maximized at a two-cluster solution (**Figure S5A**).

Individual SPNs created from covariate-residualized sulcal phenotypes had almost identical structure with SPNs created from original sulcal phenotypes (mean r = 0.996) (**Figure S4D**), suggesting that sulcal phenotypes can change with, *e.g.* age, but the relative differences between sulci remain a fixed and individualized property of the brain. Supporting this idea, the effects of standardized age, sex, and TBV effects on SPN edge weights were considerably less than their effects on the 5 MRI metrics constituting the feature vector at each sulcus (mean absolute standardized effect size: age = 0.014, sex = 0.069, TBV = 0.044, **Figure S4C, Table S3**). Given these observations, we chose to interpret SPN structure generated from the original sulcal phenotypes in subsequent analyses. Individual SPNs showed consistent topology at the node and edge levels and were highly robust against the selection of subjects or sulci included in the study (**Figure S5B-S5E**), so we next explored the linear and complex patterning of the group mean sulcal phenotype network.

### Sulci can be graded along a bipolar, linear-to-complex dimension that coheres with diverse gradients of cortical structure and function

We mapped the group mean SPN to a single, continuous dimension by computing its first principal component (PC) (**Figure 4A, Table S4**), which accounted for 27% of sulcal (co)variance across the cortex. We chose to focus on the first PC as additional PCs accounted for substantially less (co)variance, had decreased interhemispheric consistency in PC loading, had increased inter-subject variability in subject-level PC loadings (**Figure S6A**), and were more prone to outlier PC loadings (**Figure S6B**). Each sulcus was scored on the first PC, or eigen-fold index (EFI), and there was a clearly bipolar distribution of EFI scores (consistent with the two-cluster community structure), with most sulci scoring close to the linear (-1) or complex (+1) poles of the EFI and relatively few sulci having intermediate EFI scores (∼0). As shown for 10 illustrative examples, sulci with linear polar EFI scores were identifiably straighter and deeper than the more branched and fragmented sulci with complex polar EFI scores (**Figure 4A**). In these examples, it can be seen that linear sulci are dominantly composed of a single, primary sulcus, whereas more complex sulci exhibit additional secondary and tertiary branching.

**Figure 4.**
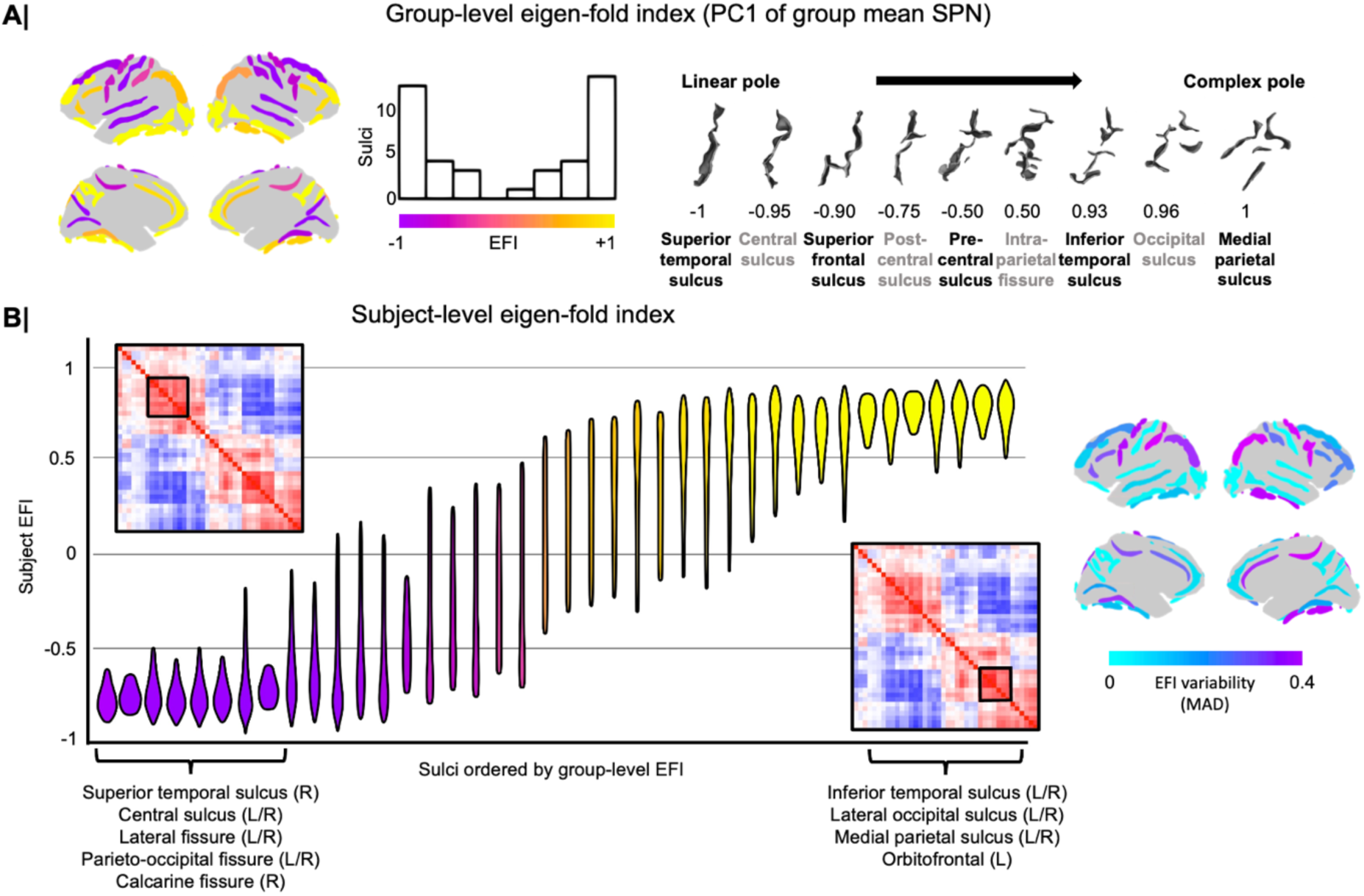
Adult sulcal patterning can be dimensionally represented by the eigen-fold index (EFI, first PC of the SPN) with both the most linear sulci (EFI ∼ -1) and the most complex sulci (EFI ∼ +1) having the lowest between-subject variability. (**A**) The EFI was derived as scaled values from PC1 of the group mean SPN, displaying a bipolar distribution with most weights for sulci at extreme ends of the dimension. Prototypical sulci were plotted in order from lowest to highest group-level EFI, depicting how this latent dimension captured linear to complex sulcal morphology. (**B**) Subject-level EFI was estimated from PC1 of subject-level SPNs, with sulcus-specific EFI distributions plotted with sulci ordered from lowest to highest group-level EFI. Violin plots displayed the middle 80% of EFI distributions for each sulcus to highlight differences in the spread of EFI values for each sulcus. Sub-clusters from the group mean SPN are highlighted and were found to correspond to the lowest and highest ranking EFI values that also had the lowest inter-subject variability. Plots of inter-subject variability in sulcal complexity as measured by EFI demonstrated that sulci with high EFI variability could be directly adjacent to sulci with low EFI variability, such as with the pre/post-central sulcus adjacent to the central sulcus.

The EFI score for each sulcus was also measured for each individual SPN, which provided a metric of inter-subject variability in the location of each sulcus on this bipolar (linear-to-complex) dimension of sulcal morphometry (**Figure 4B, Table S4;** see **Methods**). The correlation of subject EFI scores across subjects for any two sulci was low, just as with correlations from individual sulcal phenotypes (median absolute correlation = 0.04) but was generally higher for interhemispheric sulcal EFI correlations (median absolute correlation = 0.13). We found that the sulci which were closest to the linear pole (EFI score ∼ -1), and the sulci which were closest to the complex pole (EFI ∼ +1), had very low between-subject variability of EFI scores; whereas the sulci with less extreme EFI scores were markedly more variable between individual brain scans. For example, the right superior temporal sulcus, the right and left (R/L) central sulcus, R/L lateral fissure, R/L parieto-occipital fissure, and the right calcarine sulcus had the most linear polar EFI scores and had very low between-subject variability. These sulci additionally formed a sub-cluster in the linear cluster of the group mean SPN. In contrast, the R/L inferior temporal sulcus, R/L lateral occipital sulcus, R/L medial parietal sulcus, and the left orbitofrontal sulcus, formed a sub-cluster in the complex cluster of the group mean SPN, had the most complex polar EFI scores, and had very low between-subject variability. Between these poles, many other sulci varied widely between individuals in their position on the linear-to-complex spectrum of sulcal morphometry (**Figure 4B**). The greatest variability in EFI scores was seen for the left pre-central sulcus and left intra-parietal fissure - suggesting that these folds are subject to the weakest developmental constraints relative to those folds at EFI poles. This “tethering” of the EFI at highly constrained poles meant that overall organization of sulcal morphology showed a highly stereotyped topography across individuals: the cross-sulcus correlation of individual EFI scores with the group-level EFI scores was strongly positive (mean = 0.78, standard deviation = 0.09).

Given such striking conservation of SPN clusters and the EFI axis across individuals, we hypothesized that the spatial ordering of folds by EFI score or SPN cluster would cohere with other recently described organizational axes of cortical structure and function.

We first sought to validate the EFI against independently derived descriptions of cortical structure based on BrainVISA metrics not included in SPN computation including: allometric scaling of sulcal surface area^53–55^, cortical thickness of areas adjacent to sulci^56^, and mean sulcal depth heritability^23^ (**Methods**, **Figure 5A**). These analyses revealed that: (i) EFI was positively correlated with brain-size dependent expansions of cortical surface area, *i.e.*, cortical areas with more complex sulcal patterning had greatest relative surface area expansion with increasing brain size (**Figure 5A**, left); (ii) EFI was positively correlated with cortical thickness, *i.e.*, cortical areas adjacent to complex sulci were thicker than cortical areas intersected by linear sulci (**Figure 5A**, middle); and (iii) EFI was negatively correlated with an independently-generated estimate of sulcal depth heritability, *i.e.*, linear sulci had greater heritability than more complex sulci (**Figure 5A**, right; all |r|>0.5, *P*<0.009).

**Figure 5.**
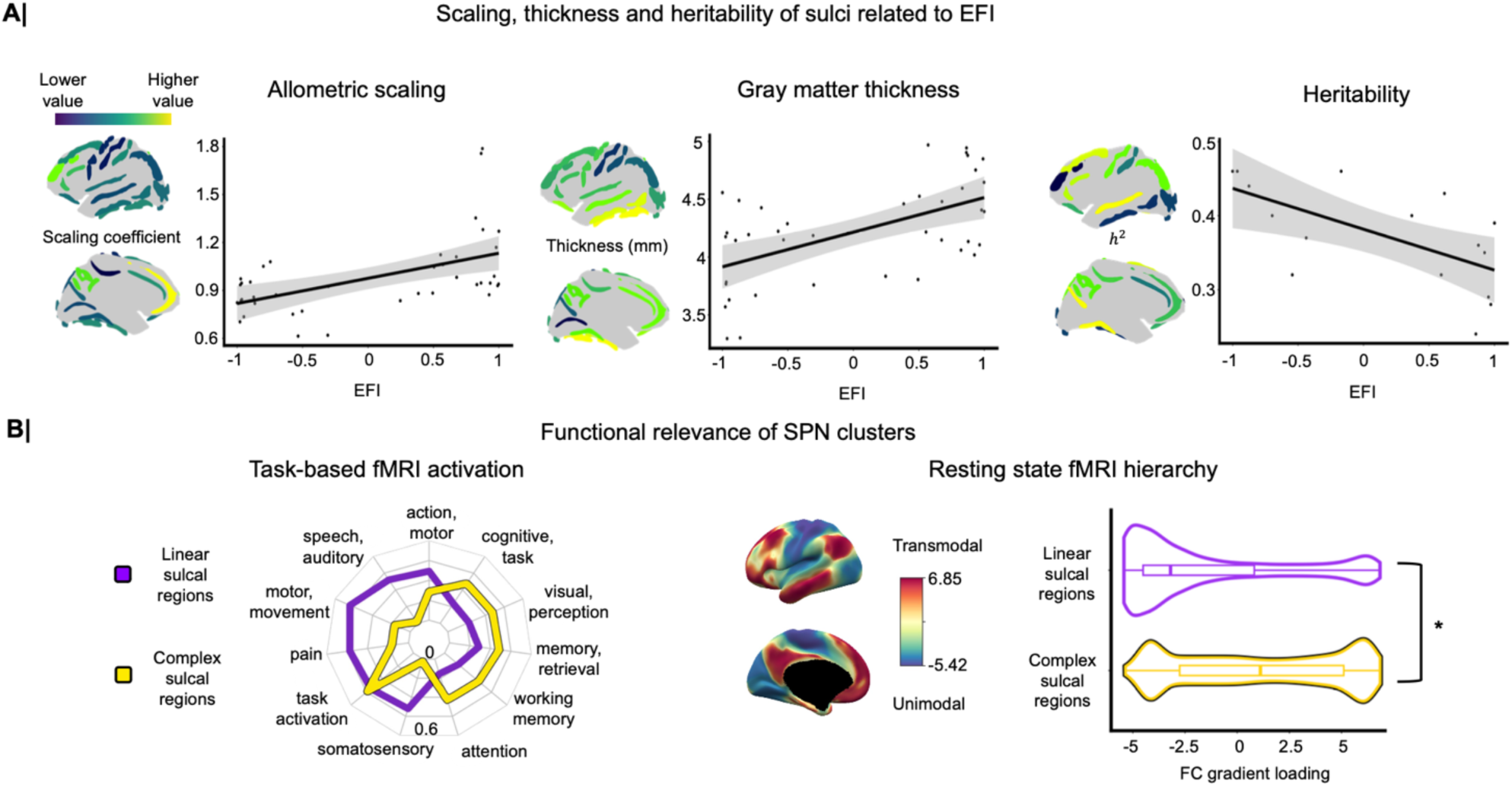
Dimensional (EFI) and categorical (SPN modular) metrics of sulcal patterning are related to cortical structure, heritability, and function. **(A)** Estimates of sulcal allometric scaling, gray matter thickness, and mean depth heritability were all derived from BrainVISA sulcal morphometry, allowing direct comparison against EFI. Heritability estimates were available for a subset of displayed sulci (**Methods**). EFI positively correlated with allometric scaling (r = 0.52, *P* = 0.00055) and gray matter thickness (r = 0.56, *P* = 0.00016) and negatively correlated with heritability (r = -0.63, *P* = 0.0086). Therefore, deeper, more linear sulci were more heritable, located in thinner cortex, and received less disproportional expansion of surface area in larger brains. **(B)** SPN clusters were convergently distinguished by task and resting state fMRI networks. Task-based functional annotations from Neurosynth revealed a sensorimotor to association bias from linear to complex sulci based on Dice overlap. Resting-state functional connectivity (FC) gradients supported the relative enrichment of linear sulci for unimodal regions and of complex sulci for transmodal regions (*P_spin_* = 0.0176).

We next compared SPN linear and complex clusters to functional MRI (fMRI) measures of cortical function during tasks and at rest (**Figure 5B**). Comparison with meta-analytic maps of task-related brain activation from over 11,000 fMRI studies (https://neurosynth.org)^57^ revealed that cortical regions adjacent to linear sulci (EFI < 0) tend to be activated by somatosensory tasks, whereas regions adjacent to complex sulci (EFI > 0) are typically activated by tasks involving higher order cognitive and association tasks (**Methods**, **Figure 5B**, left). This functional differentiation of linear vs. complex folds was echoed by comparison with the topography of functional connectivity within the cortex at rest. Specifically, cortical areas surrounding linear and complex sulci, respectively, were differentiated in terms of their loadings on the sensory-to-association cortical gradient derived from diffusion map embedding of resting state fMRI^58^. Cortical areas surrounding linear sulci had significantly lower (*P_spin_* = 0.0176) functional connectivity gradient loadings, typical of unimodal cortex, compared to the higher loadings of areas adjacent to complex sulci, typical of heteromodal cortex.

Taken together, these categorical and dimensional analyses of adult sulcal patterning overall reveal a bipolar axis from linear to complex morphology. The distinct ordering of sulci along this shape axis not only coheres with other structural and functional measures but demonstrates constraints at the most linear and complex poles, suggesting potentially different modes or phases of linear and complex sulcal development *in utero*.

### Fetal sulcation is linked to adult sulcal patterning

Based on these results and prior work^2,3^, we hypothesized that the linear-to-complex dimension of adult sulcal patterning could be rooted in the phased emergence of cortical sulci during fetal brain development. We directly tested this hypothesis by using brain MRI scans *in utero* for N=228 fetuses aged 21-36 weeks GA (**Figure 6A**) to estimate sulcus-specific trajectories or growth curves for precise quantification of key milestones in the process of sulcation for comparison with sulcus-specific EFI scores in adulthood.

**Figure 6.**
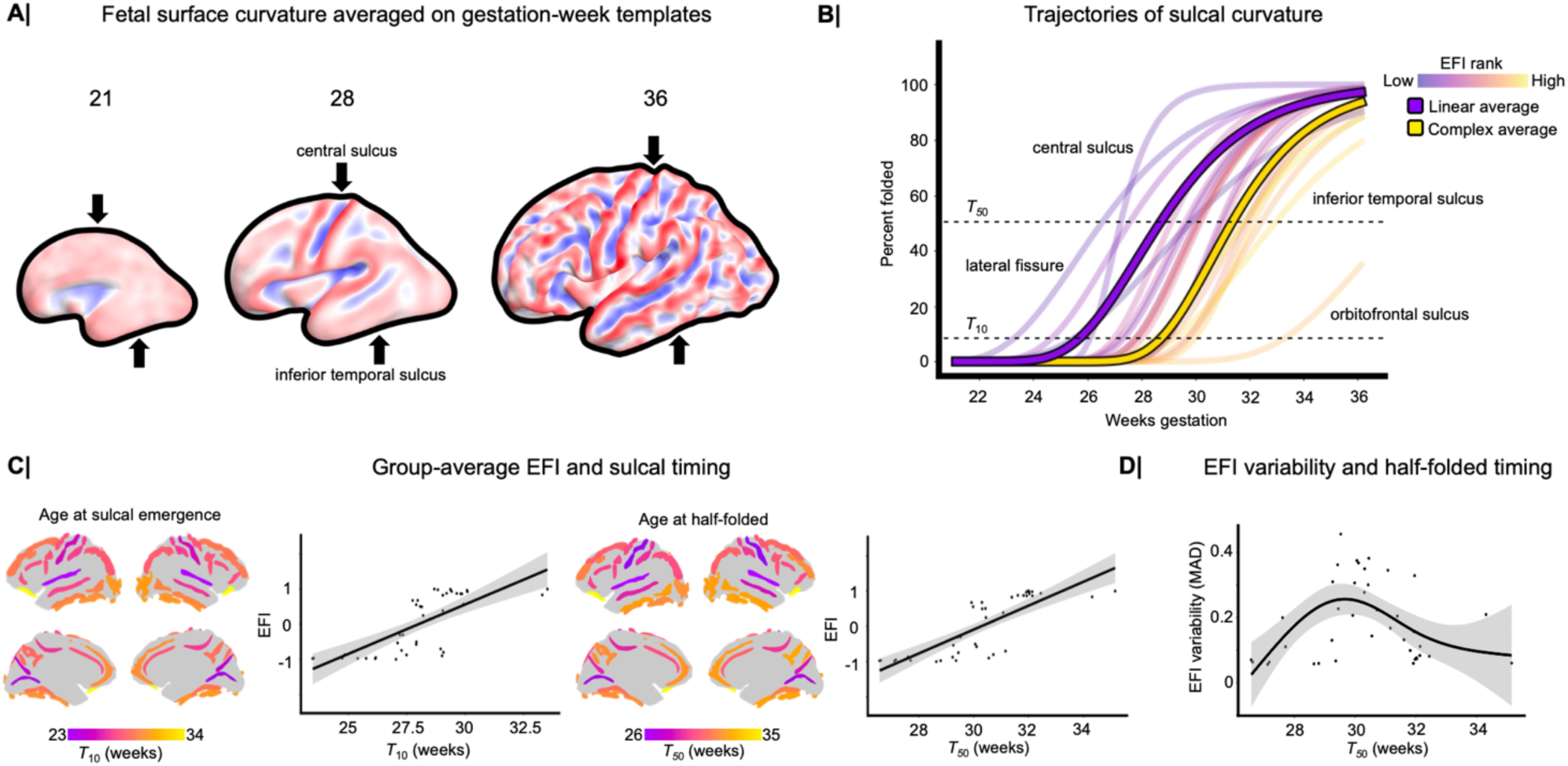
Trajectories of sulcal development in fetal brains are related to categorical (SPN modular) and dimensional (EFI) measures of adult sulcal patterning. (**A**) Exemplar cortical surface representations of the human fetal brain as it develops folds from gestational weeks 21-36, with exemplar polar linear (central sulcus) and polar complex sulci (inferior temporal sulcus) highlighted. Color codes bilaterally averaged sulcal curvature with negative (convex) curvature in red and positive (concave) curvature in blue. (**B**) Trajectories of mean sulcal curvature as a function of gestational age were expressed as a percentage of the maximum or final curvature of each (bilaterally averaged) sulcus. Bilaterally averaged trajectories are shown for visualization purposes, but hemisphere-specific trajectories were resolved and used in subsequent analyses. Individual sulci were colored in a semi-transparent scale by their EFI ranking (least to greatest). Solid colors showed trajectories for curvature averaged from either all linear or all complex sulcal regions. A sulcus was considered to have emerged once it reached 10% of maximal folding (*T_10_*), at which point approximately the sulcal indentation is first visible to the human eye; and a sulcus was designated as half-folded when it passed the threshold of 50% of maximal curvature (*T_50_*). The time-courses of sulcal emergence can also be visualized in the context of animations of temporally interpolated fetal brain MRI templates (**Supplemental Movies 1 and 2**). Linear sulci (with EFI close to -1) fold on average earlier than complex sulci (with EFI close to +1), but there is large variation in the timing of 10% or 50% maximal folding within both linear and complex clusters of sulci. (**C**) Gestational age at *T_10_* and *T_50_* was strongly correlated with sulcal differences with adult average EFI (*T*_10_: r = 0.73, *P* = 7.92 * 10^-8^; *T*_50_: r = 0.81, *P* = 3.43 * 10^-10^), indicating that more complexly branching sulci form later in development. The association is slightly stronger with *T_50_*, near when folds undergo highest rates of regional expansion. In subsequent analyses, we focused on the gestational age at half-folded. (**D**) Individual differences in sulcal EFI are related to *T_50_*, with the earliest and latest folding sulci demonstrating much less between-subject variability in adult EFI than sulci folding at intermediate gestational ages (*P* = 0.0015). Lower individual variability in EFI of the first and last developing sulci presumably represents higher genetic or geometric constraints on their shape complexity.

By tracking mean sulcal curvature over fetal life with a Gompertz growth model^59^, we captured development from 0% folding (lissencephaly, at 21 weeks GA) to 100% folded (relative plateau of curvature development, at 36 weeks GA) for each sulcus. Model fits were strong, with R-squared >80% for 95% of sulci. The models were visualized with animations to confirm they matched the visual evolution of cortical folding along spatially interpolated fetal cortical surface templates (**Supplemental Movies 1 and 2**). From these curves, we estimated key milestones in development of each sulcus, including the time to reach 10% maximum curvature, *T*_10_, and the time to reach 50% maximum curvature, or half-folded, *T*_50_ (**Figure 6B, Table S5**). We found that both milestones of fetal sulcation were strongly positively correlated with EFI scores of the corresponding sulci in the adult brain (*T*_10_: r = 0.73, *P* = 7.92 * 10^-8^; *T*_50_: r = 0.81, *P* = 3.43 * 10^-10^). In other words, sulci that had later sulcation milestones had more positive EFI scores indicating more complex adult patterning. Convergently, sulcation milestones were significantly different between linear and complex clusters of the group mean SPN, in terms of *T*_10_ (linear cluster mean = 26.6 weeks GA, complex cluster mean = 29.4 weeks GA; *t* = -5.4, *P* < 0.05) and *T*_50_ (linear cluster mean = 28.9, complex cluster mean = 31.7; *t* = -6.5, *P* < 0.05) (**Figure 6C**). The inter-subject variability of adult sulcal morphology, quantified by the median absolute deviation (MAD) of sulcal EFI scores, had a significant non-linear relationship with sulcation milestones (*P* = 0.0015), as expected from prior results (**Figure 4B**). Sulci with earliest or latest milestones of emergence had much less inter-subject variability than sulci that emerged midway through the neurodevelopmental cascade of sulco-gyral patterning (**Figure 6D**). Thus, we found that differential timing of fetal sulcation across the cortex was strongly associated with the typical form and individual differences of each sulcus on the linear-to-complex dimension of adult sulcal patterning.

### Assessment of gene expression differences across sulcal boundaries

It has previously been theorized that fetal sulcation could be driven by “buckling” of the cortical sheet at the boundaries between early-established zones of cytoarchitectonic differentiation between nascent cortical areas^10,15,60,61^. This mechanistic model was lent recent support by the finding that the orientation of sulci in adult human cortex tends to run orthogonal to axes of tangential gene expression change across the cortical sheet^41^; and these orthogonal intersections between the sulcal fundus and trans-sulcal gradients of gene expression were particularly pronounced around the central sulcus and parieto-occipital fissure. These observations suggest a mechanistic model for linear sulcation by differential cortical expansion at the boundary between cytoarchitectonically distinct cortical areas. This model would predict that (i) tangential gradients of gene expression change in adulthood should be greater in cortical zones adjacent to linear vs. complex sulci, and (ii) genes with expression gradients that are strongly orthogonal to the fundus of linear folds in adulthood should proxy early-established cytoarchitectonic differences between neighboring cortical regions. We were able to directly test these mechanistic predictions by aligning the sulcal maps provided by our automated morphometry pipeline (**Methods**) with spatially fine-grained cortical maps of tangential expression changes for 20,781 genes from Wagstyl et al.^41^

Confirming our first prediction, we found that the mean transcriptional gradient across all genes computed in Wagstyl et al.^41^ was indeed relatively increased in regions adjacent to bilaterally linear sulci, compared to regions adjacent to bilaterally complex sulci (*P_spin_* = 0.0152) (**Figure 7B**). By focusing on the principal angle and magnitude of tangential gene expression changes in the vicinity of linear sulci (**Methods**), we found that the central sulcus and parieto-occipital fissure in particular had pronounced trans-sulcal gradients of gene expression change aligned orthogonally to the fundus of each sulcus (*P_spin_* < 0.05) (**Figure 7C**). These tangential shifts in gene expression likely capture interregional differences in cytoarchitecture and are most prominently aligned with folds in regions that appear to attain their adult transcriptional identity in early prenatal life^41^.

**Figure 7.**
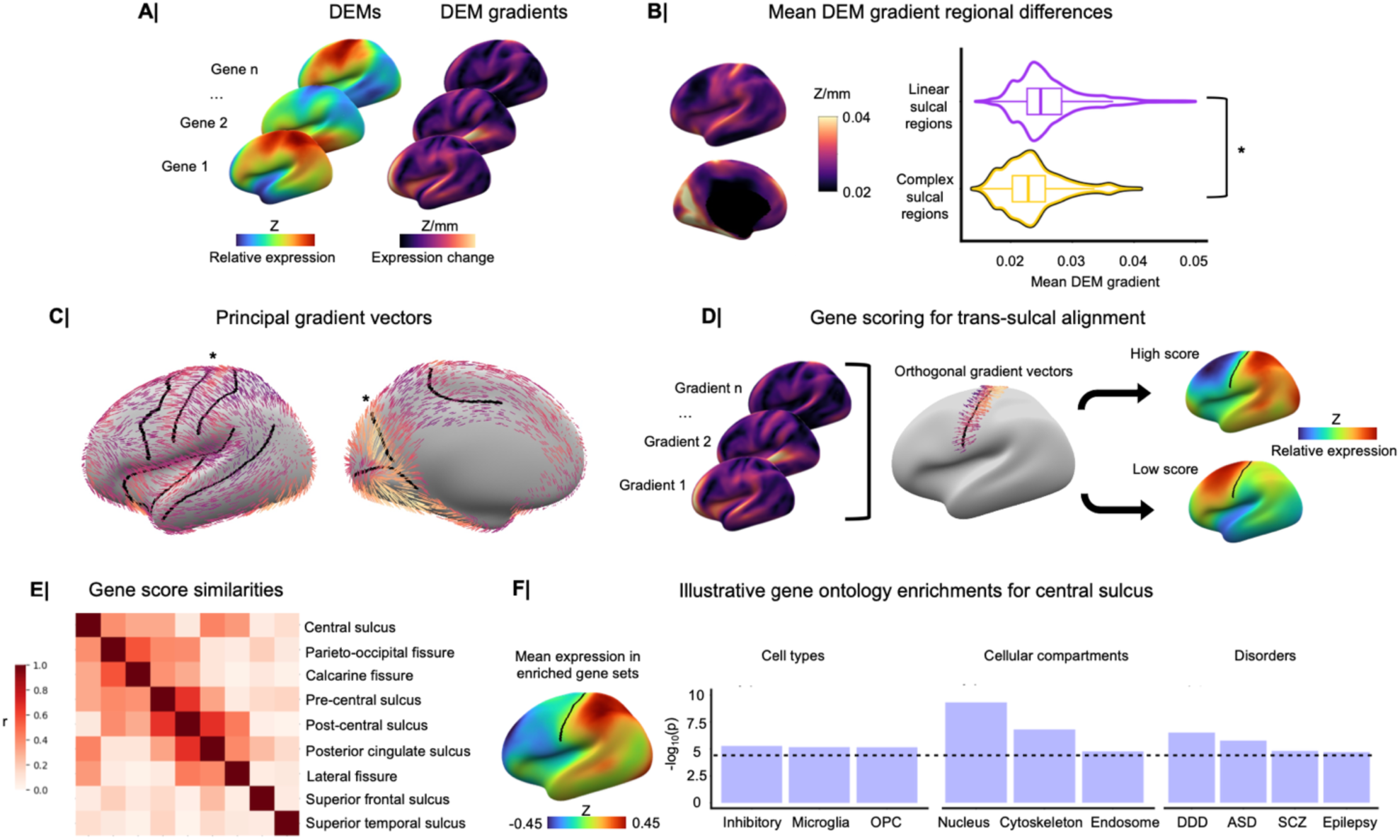
Developmentally salient cortical gene expression gradients are delineated by archetypal linear sulci. **(A)** Dense expression maps (DEMs) and their associated gradients were collated for 20,781 genes. **(B)** Linear sulci had increased average DEM gradient compared to complex sulcal regions relative to a spatial-permutation null model, indicating linear sulci were more likely to have signatures of in tangential expression change (*P_spin_* = 0.0152). **(C)** The principal gene expression vectors, or dominant directions of gene expression change, significantly aligned with the central sulcus and parieto-occipital fissure compared to random spins of sulcal fundi to random positions about a sphere, as used in subsequent analyses (*P_spin_* < 0.05). **(D)** Sulcal fundi and gene expression were mapped to a common space from which gradient vectors and fundus orientation vectors were computed. Gene scores were given by the strength of the gradient and orthogonality (angle sine) of gradient and fundus vectors, with extreme low or high scores representing strong expression changes orthogonal either direction through the sulcal fundus (*e.g.,* pre-post or post-pre central gyrus). **(E)** {1 x 20,781} vectors of gene scores per linear sulcus were correlated, with absolute values of correlations higher between proximal or parallel sulci and lower between anatomically distant sulci. **(F)** Exemplar gene ontology enrichments for gene scores are shown, with all central sulcus enrichments showing transitioning from low to high expression about pre-to post-central sulcus in the mean map across all exemplar gene sets shown. The dashed line indicates the significance threshold, and displayed P-values are capped at *P* = 1/10000.

To test our second prediction, we sought to characterize the genes that had significant trans-sulcal gradients of expression across each of 9 linear sulci (**Figure 7D, Table S6**). These gene rankings were most highly correlated between proximal or parallel sulci but were more weakly correlated between more anatomically distant sulci (**Figure 7E**), suggesting that linear sulcation is underpinned by different trans-sulcal gradients of expression in different cortical regions^62^. Gene ontology enrichment analysis of trans-sulcal expression gradients for the central sulcus and parieto-occipital fissure highlighted multiple developmentally salient gene sets (*P_spin_* < 0.05, Bonferroni corrected, **Figure 7F, Table S7**). For example, gene expression gradients orthogonal to the central sulcus were enriched for inhibitory neurons, microglia, oligodendrocyte precursor cells, cytoskeleton compartment, nuclear compartment, endosome compartment, and rare risk genes for autism spectrum disorder, schizophrenia, severe developmental disorders, and epilepsy. Each of these gradients represented a transition from low expression in pre-central gyrus to higher expression in post-central gyrus. The trans-sulcal gradient for the parieto-occipital fissure was distinctively enriched for layer V marker genes with expression generally increasing superior to the fissure in medial parietal cortex. Additionally, since adult and fetal cell marker genes for cytoarchitecture share patterning in dense expression maps (DEMs)^41^, we further tested for enrichment of fetal cell types. Fetal cell enrichments for the central sulcus were oriented along the same pre-to-post-central gyrus direction and included excitatory neurons (ExDp1/ExM-U), inhibitory migrating neurons (InMGE/InCGE), and intermediate and cyclic progenitor cells (PgG2M/PgS). Taken together, these findings confirmed our second hypothetical prediction that the linear sulci in the adult brain are orientated such that they intersect or bisect large zones of rapid tangential change in gene expression.

## Discussion

We applied innovative tools for sulcal morphometry to large-scale MRI datasets on adult and fetal human brain structure to discover a bipolar organizational axis of sulcal phenotype networks linked to the timing of sulcal emergence *in utero*. By incorporating complementary information from diverse datasets, including measures of cortical structure, function, and gene expression, we provide a deep annotation of this new perspective on human cortical surface patterning. We release this annotation in detail, along with containerized tools for automated sulcal morphometry and SPN analysis, to accelerate future research on cortical folding. We consider each of the main outputs of our work in more detail below.

### A new taxonomy of adult sulcal morphology

Many sulco-gyral analyses have focused on fold typology^7^, namely in assigning the sulcal pattern of a given region to one of multiple categories. Such approaches have been successful in parsing some variance in psychiatric disorder based on which fold types are present, but pattern types are only well-defined for a subset of cortical folds^14^. Therefore, we chose to focus on sulcal properties inherent to all sulci, *e.g.*, longest branch length, that underlie differences in sulcal typology, *e.g.*, connected or continuous cortical folding patterns of the anterior cingulate cortex^6^ or orbitofrontal cortex^63^.

High-throughput sulcal morphometry in the UK Biobank cohort provided a unique dataset for quantitative analysis of human sulcal variation. Our approach integrated complementary diverse metrics of sulcal shape that have traditionally been considered in isolation from each other. For example, sulcal depth has been related to cognition and developmental disorder^3,13,28,29,49^. Depth variability^52^ captures sulcal pits and *plis de passage* (respectively, local minima and maxima of depth along the sulcal fundus) which are genetically constrained and functionally important^64,65^. Multiple studies have examined sulcal length in relation to developmental disorders and genetics^23,27,29,53^, and cortex-wide differences in the length of the longest branch of each sulcus were noted in Chi et al.^3^ and Ono et al.^14^ Auzias et al.^66^ found that the sulci with longest branches were sufficiently consistent in their orientation to define latitudinal and longitudinal axes of the cortex. Only a few other studies have explicitly considered fractal geometry of brain gyrification^67^ or sulcal shape, including studies of the fractal dimension (FD) of sulcal surfaces embedded in a 3D volume^48,50^, with 2 < FD < 3. Here we used different but related fractal measures, focused on the embedding of sulcal lines tangent to the cortex, with ∼1 < FD < 2, as innovative measures of sulcal non-linearity or complexity (**Figure 1**).

Leveraging this diverse set of sulcal shape phenotypes, we revealed two archetypes of sulcal organization that were convergently represented by distinct clusters of the sulcal phenotype network (SPN) and by the bipolar distribution of sulci on a linear-to-complex dimension (EFI). Linear polar sulci, *e.g.*, central sulcus and parieto-occipital fissure, were typically deep, straight (FD∼1) indentations across unimodal cortex and were more heritable than complex polar sulci, *e.g.*, orbito-frontal sulcus, which were typically shallow, complex (FD > 1) and located in heteromodal cortex. This bipolar taxonomy of sulcal patterning cohered with diverse other axes of cortical organization including allometric scaling, gray matter thickness, and sensorimotor-association functional hierarchy^68^ (**Figure 5**).

### Timing of fetal formation of sulci was linked to polarity of adult sulcal shape

The clear bipartite clustering of the group mean SPN (**Figure 3B**) aligned approximately with prior measurements of the timing of sulcal formation during fetal brain development. For example, most of the sulci in the linear cluster were first visible postmortem in brains less than or equal to 25 weeks GA, and all of the sulci in the complex cluster were not discernible until later in fetal development^3^. However, we wanted to explore the developmental provenance of adult sulcal patterning more directly and quantitatively. To do this, we reconstructed the cortical surfaces for 228 fetal brain MRI scans and estimated high temporal resolution trajectories of curvature development for each of 40 sulci identifiable in adult brains. We were then able to link the timing of fetal sulcation to the polarity of adult sulcal shape, comprehensively and based on a much larger fetal brain dataset than has been reported in postmortem studies.

We confirmed that linear sulci (with EFI ∼ -1) formed earlier than complex sulci (with EFI ∼ +1), *e.g.*, there was a strongly positive correlation between adult sulcal EFI and the gestational age at which that sulcus was 50% (or 10%) fully formed *in utero* (**Figure 6**). Slightly stronger correlation was found with the 50% (*T_50_*) milestone (r = 0.81), possibly because this milestone marks the time point at the greatest rate of sulcal curvature development, likely co-occurring with rapid neurodevelopmental programs of tangential expansion of the cortical plate^69^. We also confirmed prior expectations that between-subject variability or individual uniqueness of adult sulcal shape should be minimized for linear sulci that formed earliest^7,22^.

Knowing that the earliest-forming, most linear sulci are more heritable, it is intuitive that their development should be stereotypically determined by a genetic program that is expressed regardless of stochastic or environmental differences between individual brains. However, more counter-intuitively, we also observed low levels of between-subject variability of adult shape for the most complex, latest-forming sulci (**Figure 4**), *i.e.*, there was a non-linear, inverted-U relationship between individual variability of sulcal EFI and the milestone of 50% complete fetal sulcation (**Figure 6**). The most individually variable sulci had intermediate, less polarized values of EFI on average; emerged at intermediate gestational ages between the earliest, linear and latest, complex sulci; and were sometimes directly proximal to sulci with the lowest between-subject variability of EFI (**Figure 4**). This variability could reflect competing mechanisms of cortical arealization driven by proximity to primary or unimodal cortex^70^ met with stochastic buckling pressures midway through sulcal ontogenesis. For example, small inhomogeneities in initial cortical geometry or tissue properties have been found to have substantial consequences on final sulco-gyral patterning^15,71^.

These observations suggest that fetal sulcation is more strongly constrained (less individually different) at the start and the end than in the middle of the cascade of cortical folding. But the nature of such constraints remains to be parsed. Some studies frame sulcal development as a homogenous process wherein all sulci form by the same mechanisms^10^, but recent models have begun to shed light on mechanistic heterogeneity between different sulci^69^. As noted, genetic constraints could explain the reduced sulcal complexity variability for the first-forming sulci. However, their relatively low heritability suggests that non-genetic constraints must apply to the latest forming, complex sulci. Just as models of crumpled paper revealed constraints on how further crumpling is constrained to maximally relieve tension in material^72^, the final cortical folds to form *in utero* could experience physical constraints on their formation in the context of multiple prior sulci having already “carved up” the cortical sheet available for new sulcation. The prior presence of sulci very likely impacts the later formation of proximal sulci^15,71^, and it is an important question for future studies to resolve more precisely how the end-point of the fetal sulcation cascade could be physically constrained.

### Trans-sulcal expression gradients as a mechanism for linear sulcation

One prominent mechanistic model for sulcation is that the cortical sheet buckles under the physical stress of differential tangential expansion of neighboring but distinct cortical areas, or between different layers of the same cortical area. In differential tangential expansion (DTE) models, cortical surface gyrification results from a more rapid expansion of a stiffer cortical plate relative to deeper fetal tissue compartments within the same area^69^. Elaborations of DTE models have included differential expansion across regions in the cortical plate^71,73^. Cortical differentiation of cytoarchitectonically well-defined areas could lead to predictable location of sulcal infolding and it is compatible with this model that some sulci have long been recognized to demarcate cytoarchitectonically distinct areas^60^; *e.g*., the central sulcus and parieto-occipital fissure both separate clearly distinct areas. More recent histological work^61^ demonstrated that primary fissures, aligning with our notion of linear sulci, were better predictors of cytoarchitectonic boundary than sulci in higher order or associative cortex, aligning with more complex sulci.

By this account, the topographical patterning of cortical gene expression that controls areal differentiation should be related to linear sulcal anatomy, with differential gene expression either side of the sulcal line or fundus being compatible with differential expansion of the adjacent cortical areas divided by the fundus. To test this model, we used dense expression maps (DEMs) of cortical gene transcription that allowed us to measure the magnitude and geometric alignment of tangential expression gradients in the vicinity of specific sulcal fundi. First, as predicted hypothetically and by prior work^60,61^, we found that the magnitude of tangential gene expression gradients was significantly greater in cortical regions adjacent to linear sulci than in regions adjacent to complex sulci, indicating that linear sulci were indeed co-located with more locally variable patterns of cortical gene expression. Second, we focused on a more detailed analysis of the angle as well as the magnitude of gene expression gradients with respect to 9 linear sulci, including the central sulcus and the parieto-occipital fissure. We reasoned that gene expression gradients driving sulcus formation by differentiation of adjacent cortical areas should be orientated orthogonal to the sulcal fundus. We identified and functionally annotated gene sets that had such significant trans-sulcal gradients of expression for each sulcus individually. We found there was some consistency between physically proximal sulci in terms of their trans-sulcal expression gradients, but also clear evidence that different sulci were associated with genetically and functionally distinct trans-sulcal gradients of expression. For example, the trans-sulcal gradients that cut across the central sulcus were enriched for genes associated with neuronal development and neurodevelopmental disorders, whereas the genes that were most differentially expressed either side of the parieto-occipital fissure were enriched for layer V neurons of medial parietal cortex.

These coordinated configurations of folding and gene expression in adulthood are consistent with a biophysical model where early-established cytoarchitectural gradients predispose the buckling cortical sheet to form sulci in stereotyped locations. It is also possible - although less likely - that the stereotyped location of linear sulci is aligned with but not caused by cytoarchitectonic differences, or is itself a mechanism by which cytoarchitectonic differences are achieved between neighboring cortical regions^10,74^. It will require further investigation to validate these trans-sulcal expression gradients as mechanisms for linear sulcation but our observations are consistent with this hypothesis in general and suggest that specific linear sulci may be formed by specific gradients of gene expression related to the relative differentiation of the cortical areas they divide.

### Tools for high throughput sulcal morphometry and SPN analysis

To facilitate future investigations, we provide our suite of new metrics for comprehensive characterization of sulcal morphology, which drove our main discovery of a bipolar taxonomy convergently defined by clustering of sulcal phenotype networks and a linear-to-complex dimension of sulcal patterning. The 5 morphological metrics of average (median) sulcal depth, depth variability (median absolute deviation), longest branch, branch span, and fractal dimension, can be computed within ∼2 minutes for each MRI scan processed with BrainVISA Morphologist. Because FreeSurfer software improves BrainVISA’s sulcal segmentations^29^ and has its pre-processed outputs often included in neuroimaging dataset releases, we developed a containerized pipeline which generates full sulcal morphometrics, SPN and EFI results from FreeSurfer outputs for a single scan in ∼25 minutes, *e.g.*, on a single CPU, 8 GB RAM, Linux machine, from a single command.

### Limitations and future considerations

There are several limitations inherent to the datasets and approaches used in this study. First, while the large UK Biobank cohort of adults encompassed the generally high levels of between-subject variability in sulcal patterning, such variability could be partly driven by maturational and age-related changes not directly linked to generative mechanisms in fetal development. It will be important to apply identical sulcal morphometry pipelines to younger cohorts to test the generalizability of this bipolar taxonomy of sulcal patterning, and its links to fetal sulcation in younger and more diverse cohorts. Additionally, while we used the largest fetal brain imaging cohort available, questions remain regarding between-subject variability in sulcal emergence and in sulcal complexity that would only be accessible in a longitudinally scanned cohort followed up through term birth. Within-subject tracking could help overcome limitations in registering fetal cortical surfaces with limited sulcal depth patterning near ∼21 weeks GA, which could lead to noise in estimating boundaries of prospective sulcal regions.

Overall, more work needs to be done to evaluate mechanisms of cellular differentiation and differential gene expression across linear and complex sulci. However, popular model organisms for gyrencephalic development, like the ferret^18^, only have linear sulci, which could impede translational studies of the bipolar sulcal taxonomy evident in humans. To date, there are no human datasets of fetal gene expression encompassing the full 21-36 weeks GA period over which both linear and complex sulci form. And, the extant fetal gene expression data for the ∼23 week GA period^75^ do not have sufficiently fine-grained spatial resolution to measure gene expression gradients as precisely as we were able to measure them in dense expression maps of the adult brain. Even in densely sampled adult postmortem tissue, the smaller gradients of gene expression hypothetically associated with complex sulci could be difficult to detect due to reliance on registration of these highly variable sulci to landmarks in a common space for analysis^76,77^.

## Conclusion

We have introduced and applied new methods for measuring sulcal phenotype networks from MRI scans of the human cortical surface. We have presented evidence in support of a bipolar taxonomy for adult sulcal shape that is directly linked to fetal development, and in support of the more mechanistic hypothesis that linear sulcation is driven by trans-sulcal gradients of gene expression. The annotated datasets and computational tools used to generate these results are published as an open resource to facilitate future mechanistic, developmental, and clinical studies of sulcal patterning.

## Methods

### UK Biobank cohort

The UK Biobank is an effort led to collect diverse phenotypic data to promote population-level assessments of lifestyle, environment, and genetics on biology and health presentation. A subset of subjects enrolled in the study participated in brain imaging, from which the first imaging session’s data were retrieved (https://www.ukbiobank.ac.uk)^37,38^.

Scanner acquisition of brain structural MRI is detailed elsewhere (https://biobank.ctsu.ox.ac.uk/crystal/crystal/docs/brain_mri.pdf). In brief, all three scanning sites used standard Siemens Skyra 3T scanners with a Siemens 32-channel RF receive head coil. T1 and T2-FLAIR acquisitions were both downloaded to support gray and white matter tissue segmentation. T1 acquisition involved a five-minute 3D MPRAGE session at 1×1×1 mm resolution, in-plane acceleration iPAT=2, and prescan-normalization. T2-FLAIR acquisition involved a six-minute 3D SPACE session at 1.05×1×1 mm resolution, in-plane acceleration iPAT=2, partial Fourier = 7/8, fat saturation, elliptical k-space scanning, and pre-scan normalization^78^.

### Extraction of sulci and quality control

Cortical surfaces were first reconstructed for all subjects with T1 images and additionally with T2-FLAIR images if available as in Mallard et al.^79^ using FreeSurfer neuroimaging software (version 6.0.1)^44,45^ packaged in fmriprep 21.0.2^80^. FreeSurfer segmented boundaries of gray and white matter from the images, which were input to BrainVISA neuroimaging software (version compiled on August 8^th^, 2022) for watershed algorithm detection of sulcal boundaries. BrainVISA’s Morphologist pipeline was used to extract sulci as the regions filling between gyral peaks, subsequently skeletonized to the wall of voxels halfway between bordering gyri^24–26^. All sulcal phenotypes were derived from points defined on the tops and bottoms of the voxel medial walls (**Figure 1A**, right).

After excluding N = 1,568 subjects with neurological conditions^46^, three quality control measures were implemented. First, previous work has demonstrated that robust automated quality control of MRI can be performed by detecting outliers of FreeSurfer mesh reconstruction quality^81^. As is standard^82^, outlier detection of Euler Number (EN) was performed within each scanning site, with interquartile range outliers excluded from analysis (N = 2,026). The second quality control measure was to assure the accuracy of BrainVISA’s Morphologist pipeline sulcal labeling. Previous works have excluded small sulci not labeled by BrainVISA in the majority of subjects^23,27,29^. We found that retaining 51 out of 123 possible bilateral sulcal labels allowed 95% of the original sample to be retained. From this set, we excluded sulci without anatomical meaning consistent with other sulci (*i.e.*, the anterior lateral fissure label does not trace between gyri) and then merged anatomically adjacent labels (*e.g.*, merging five labels of the pre-central sulcus into one label), mitigating BrainVISA Morphologist mislabeling that typically propagates amongst anatomically proximal regions (**Figure S2**). We retained labels for 20 sulci per hemisphere that spanned the lateral, medial, and ventral faces of the cortex (**Figure S2**). A small set of subjects that did not receive a label for one of these 40 major sulcal regions were excluded from analysis (N = 191). A final, necessary quality control measure was discovered during analyses. A small set of subjects (N = 30) had outlier residuals from log-log regression models^53–55^ fit between total sulcal surface area (summed sulcal surface area from all 40 sulci) and total brain volume (FreeSurfer’s “BrainSegVol”).These subjects were visually confirmed to have been segmented poorly by FreeSurfer, leading to overestimation of sulcal surface area by BrainVISA. These subjects were also excluded from further analysis, leaving a final 34,725 subjects for analysis.

### Sulcal phenotype networks (SPNs)

SPNs were defined as the {40 x 40} matrix of correlations from {1 x 5} vectors of sulcal phenotypes between sulci in a given brain. First, sulcal phenotypes were extracted from subject scans in order to generate SPNs. Extracted sulcal medial walls contained points closest to the depths of sulci (bottom points, or fundus points) and points closest to the exterior of the brain, termed hull-junction points in BrainVISA as they lie along the hypothetical convex hull (inflated or unfolded surface) of the cortex (**Figure 1A**). Depth profiles or histograms (**Figure 1B**) were generated for any sulcus by finding the shortest geodesic path along the sulcal medial wall from fundus points to hull-junction points. Average depth was given by the median distance and depth variability was given by the median absolute deviation of distances, as in Klein et al.^52^

Exterior or hull-junction points were useful for branching pattern measures (longest branch, branch span, fractal dimension) as the full tangential sprawl of the sulcus is exposed at this minimum depth on the sulcal medial wall. Similar to length calculations native in BrainVISA, longest branch was defined as the longest contiguous geodesic path along hull-junction points given the same sulcal label. Branch span was inspired by analyses of dendritic morphology that used a “circularity index” to assess how uniform or dispersed the orientation of dendrites were about their soma^51^. This calculation is performed in a two-dimensional reference frame, so hull-junction points from contiguous sulcal branches were projected to a plane tangent to the hull of the cortex centered at the center of mass of a given sulcal branch. The sulcal branch point sets in the plane were all aligned to the center of mass of the largest sulcal branch if multiple sulcal branches were present. Branch span was given by the ratio of the convex hull area of this point set divided by its circumscribed area, as in Levy et al.^51^ Circumscribed area was derived from the smallest enclosing circle of the point set (https://www.nayuki.io/page/smallest-enclosing-circle). Finally, fractal dimension was calculated from a three-dimensional box-counting algorithm (https://github.com/ChatzigeorgiouGroup/FractalDimension) applied to the hull-junction points all assigned a given sulcal label to assess self-similarity of sulcal branching patterns on the exterior of the cortex.

Age, sex, and total brain volume (TBV) effects on sulcal phenotypes were modeled with multiple linear regression. Plots of each sulcal phenotype versus age and TBV were visually inspected and confirmed to have either linear or no relationship. Therefore, for consistency across the 200 linear models (40 sulci * 5 phenotypes), the same multiple linear regression model was used with linear terms for covariates. Bonferroni correction was performed for 120 tests (40 sulci * 3 covariates) to investigate significance within each model on a phenotype-by-phenotype basis.

All five sulcal phenotypes were calculated for all 40 bilateral sulci and for each subject to enable SPN generation. Each sulcal phenotype was Z-scored within each brain to capture within-brain inter-regional differences in sulcal phenotypes. Then, a {40 x 40} correlation matrix or SPN was generated by the sulcus pairwise correlation of the five sulcal phenotypes. Age, sex, and TBV effects on the 780 unique edges in the symmetric SPNs were modeled with multiple linear regression in the same manner as with individual sulcal phenotypes. Bonferroni correction was performed for 2,340 tests (780 edges by 3 covariates) to investigate significance within each model on an edge-by-edge basis.

### Categorical and dimensional analyses of SPNs

Hierarchical clustering of group mean SPNs was performed in R (https://www.r-project.org/), finding an optimal split of the resulting dendrogram at two clusters, given by the number of clusters that maximized the Dunn index over 2 through 10-cluster solutions. The group-level eigen-fold index (EFI) was given by the first principal component of the full group mean SPN correlation matrix, assigning a PC loading to each sulcus, specifying low (negative) EFI scores to include linearly shaped sulci and high (positive) EFI scores to include more complex sulci. Group-level EFI was scaled from -1 to 1 for interpretability. The interpretation of linear and complex sulci was determined from viewing prototypical sulci across the PC loading space and by looking at sulcal phenotype distribution by sulcal cluster. For subject-level PCA decompositions, PC signs are arbitrary, so it would not be feasible to inspect and interpret all subject-level components. Additionally, subject-level PCs could each represent slightly different organizational axes. Therefore, subject-level EFI was calculated as sulcal coherence with the group-level EFI, calculated as the correlation between a subject SPN row for a given sulcus with the EFI; see also **Figure S6A** for supplemental results on PCs beyond PC1.

### Structural and functional brain map annotation against SPNs

Structural and functional maps comparable with EFI and SPN clusters were either derived from other studies or computed in the present study. Allometric scaling coefficients represented the degree to which two-dimensional sulcal surface area scaled with three-dimensional TBV, with values above 0.67 indicating positive allometric scaling (*i.e.*, more sulcal surface area than expected given TBV). Sulcal surface area was measured as in Fish et al.^53^ for each sulcus, calculated as sulcus mean depth * length. Log-log regression models^53–55^ between age and sex residualized sulcal surface area and TBV (FreeSurfer’s “BrainSegVol”, encasing volume of the cortex) were fit and had coefficients associated with total brain volume pulled for Pearson correlation with EFI. The coefficients represented the degree to which sulcal surface area disproportionately increased with total brain volume increases.

Gray matter thickness measures for each sulcus in subject were computed by BrainVISA Morphologist, calculating thickness as geodesic distances from white to gray matter on either side of the sulcus in volumetric space^56^. Gray matter thickness values were averaged across subjects to allow Pearson correlation with EFI.

Heritability estimates were derived from mega-analytic family-based studies using BrainVISA Morphologist to measure mean sulcal depth^23^, the measure most linked to the EFI in our study. Where heritability estimates were given for multiple sulcal labels that were merged into one label in our study, we used the maximum heritability of these sub-labeled sulci, since maximum values represented the most sensitive and meaningful sub-labeled sulci. Using the mean heritability of sub-labeled sulci did not considerably change significance or interpretation of the Pearson correlation between mapped heritability values and EFI.

Neurosynth meta-analytic functional annotation of SPNs, as performed in Wagstyl et al.^41^, used the Dice overlap between functional regions and regions of interest to infer functions subserved by the regions of interest. We generated a parcellation of sulcal regions in FreeSurfer’s FreeView application (http://surfer.nmr.mgh.harvard.edu/) consistent with BrainVISA standard sulcal labeling^26^ and endpoints of sulcal regions in the Destrieux atlas^83^ to test for overlap between linear or complex sulcal clusters with functional regions. All sulcal parcellations used in this study are provided (link available prior to publication). 30 functional regions were computed from 30 topic-modeling derived terms from over 11,000 functional MRI tasks (https://neurosynth.org)^57,84^.

Significant functional regions were downloaded (https://neurosynth.org/analyses/topics/v4-topics-50/), mapped to closest vertices on an average pial surface mesh, and tested for dice overlap with linear and complex sulcal regions.

Functional hierarchy was measured as the first principal component loadings of functional connectivity from Margulies et al.^58^ and downloaded using NeuroMaps^85^. The difference in loadings between linear and complex sulcal regions was compared to a null distribution based on permuted linear and complex sulcal labels, asking whether this difference was more extreme than in randomly constructed SPN cluster definitions. Permuted labels were generated from spinning the original labels projected onto a sphere and reassigning labels to sulcal regions 10000 times, using the *gen_spinsamples* python function from the *netneurotools* package with the Hungarian algorithm selected^86^.

### Fetal cortical surface processing

Sulcal curvature was measured in each of 40 bilateral sulci for N = 228 subjects aged 21-36 weeks GA from the Developing Human Connectome Project’s (dHCP) fetal imaging cohort. Fetal MRI were passed through the dHCP structural processing pipeline^47^, with initial tissue segmentation performed by DRAW-EM^87^. Quality control of T2-weighted images, segmentations, and surface reconstructions informed subsequent manual editing of tissue segmentations by expert (V.K.) for every subject. Edited segmentations were again passed through surface reconstruction to yield final surfaces^88^. Subjects with low quality assessment scores (N = 5) from expert evaluation were visually inspected to confirm poor quality and were excluded.

Coordinates of subject surfaces were aligned with nearest age templates created for surfaces 21-36 weeks GA^89^. Surface alignment was performed using spherical representations of cortical surfaces, aligning dHCP subject maps of sulcal depth with template sulcal depth using Multimodal Surface Matching (MSM)^76,77^. Aligning surfaces via sulcal depth to nearest-week spatiotemporal templates^90^ has been found to be effective for fetal surface registration^91^. By computing vertex-wise averaging of curvature across subjects mapped to a given GA week template, we were able to visually inspect and confirm that MSM successfully aligned all major sulcal landmarks as expected (**Figure 6A**). Sulcal borders were delineated on the 36 weeks GA template in *FreeVie*w in alignment with BrainVISA standard sulcal labeling^26^ and endpoints of sulcal regions in the Destrieux atlas^83^ to allow comparison between BrainVISA sulcal measures from adults. These borders were then mapped to each next earliest GA template successively using MSM and were visually assessed for successful registration. As in Xu et al.^59^, the gray-white matter interface was used for measurement of cortical curvature^40^. Absolute values of the curvature in each sulcal region measured sulcation as the degree of deviation from a flat surface.

Following Xu et al.^40^, curvature along the grey-white matter interface surface can be cross-sectionally modeled along development using a Gompertz function. For all 40 bilateral sulci, we fit a Gompertz function using Python’s *curve_fit* function as part of the SciPy *optimize* module. All models had an R-squared of approximately 0.8 or higher, except for two sulci (left intermediate frontal and right orbitofrontal) which were visually confirmed to have models that accurately portrayed sulcal emergence. Additionally, we visually compared each model against an animation (https://github.com/StuartJO/BrainSurfaceAnimation) created from temporal interpolation of template surfaces, again confirming that the models behaved in agreement with visually recognizable timing of sulcation (**Supplemental Movies 1 and 2**). Gestational age at fitted values for 10 and 50% of the estimated plateau of Gompertz functions defined *T*_10_ and *T*_50_ sulcal development milestones, which were subject to Pearson correlation against EFI. Additionally, a general additive model with 5 knots (optimized to balance over and under-fitting) was fit between *T*_50_ and the median absolute deviation of subject EFI scores for each sulcus using the *gam* function from the *mgcv* library in R^92^.

### Transcriptional annotation of SPNs

We used the dense expression maps (DEMs) created in Wagstyl et al.^41^ to test for sulcal alignment with gene expression gradients. Microarray gene expression for each of 20,781 genes was assessed in multiple different cortical locations from six postmortem donor brains from the Allen Human Brain Atlas^75^. The spatially smoothed and averaged maps were found to replicate other known cytoarchitectonic boundaries in Wagstyl et al.^41^ In the present work, we first asked whether gradients of gene expression maps were greater in linear as compared to complex sulci, hypothesizing that gene expression would undergo more drastic changes about linear sulci. We again used spin-based permutations^86^ with 1000 spins to evaluate evidence for greater gradient magnitude in linear sulci.

Post-hoc analysis tested whether specific linear sulcal fundi marked the distinct boundary between high and low cortical gene expression. Sulcal fundi can be determined for linear sulci in average mesh representations of the cortex due to their low inter-subject variability, so we first extracted linear sulcal fundi using Automated Brain Line Extraction (ABLE)^93^. Resulting sulcal fundi lines were corrected for small errors by redrawing on the cortical surface using Connectome Workbench’s image viewer^94^. The fundi represented the midline between gyral crests that traced through the points of highest curvature at the deepest points of sulci. Sulcal fundi and DEMs were both projected to spherical representation of the cortical surface, with gene expression gradients and sulcal fundus orientation recomputed on the sphere using the Workbench *-metric-gifti* command^94^. Principal expression vectors were calculated as the first principal component of gene expression gradient vectors at each vertex across all genes. To first test whether these vectors ran orthogonal to sulcal fundi at large magnitude, each sulcal fundus was scored with the following equation:

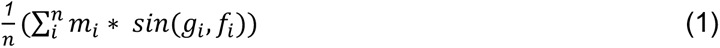

where n = number of vertices in a sulcal region, *m_i_* = magnitude of the mean gradient at vertex *i*, *i* = *i*^th^ vertex, *g_i_*= gradient vector at vertex *i*, and *f_i_* = fundus orientation vector for the closest fundus vertex to vertex *i*. For each sulcus, this value was recomputed following 10,000 random spins of the sulcal fundus about the sphere, ensuring the spun fundus did not cross into subcortical vertices. The degree to which all vectors pointed the same direction was not considered as the sign of the principal expression direction is arbitrary. For gene-level scores for each sulcus, the same score was used but also had to consider the degree to which all vectors pointed the same direction about the sulcal fundus, given by the equation:

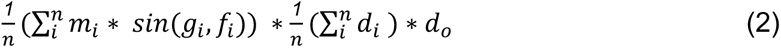

where *d_i_* was a binary variable representing the direction the gradient vector pointed to (*e.g.*, 0 = left-to-right through sulcal fundus and 1 = right-to-left through sulcal fundus when the majority of vertices had gradient vectors oriented right-to-left) and *d*_0_(overall direction) was -1 or 1 depending on the direction the majority of vertices’ gradient vectors pointed through the sulcal fundus. This calculated the strength of orthogonal gene expression gradient to sulcal fundi, weighted by the angle, amount of vectors pointing the same direction, and magnitude of gradient. Thus, signed gene scores gave information on the direction and coherence of the trans-sulcal alignment of gradient vectors with the sulcal fundus. Gene ontology enrichment for gene scores were computed using spatial permutation nulls, wherein the sulcal fundus was randomly rotated 10,000 times with gene scores recomputed for each rotation. As in Fulcher et al.^95^, the mean gene score for a gene category was tested for whether it was more extreme than mean gene scores from spatially permuted data. Bonferroni correction accounted for two sulci (significant from principal expression analysis) times two directions from which genes could transition from low to high or high to low expression.

## Data and code availability

Code for the derivation of sulcal phenotypes and SPN analysis is made available (link available prior to publication). We provide this code as a containerized pipeline (available prior to publication) to derive sulcal phenotypes directly from brain scans processed by the frequently used FreeSurfer neuroimaging software.

## Acknowledgments

W.E.S. is a PhD student/candidate in the NIH Oxford-Cambridge Scholars Program and is also supported by the Gates-Cambridge Scholarship. W.E.S. and A.R. are supported by the Intramural Research Program of the National Institute of Mental Health (NIH annual report number ZIAMH002949). This work received support from computational resources of the NIH HPC Biowulf cluster (http://hpc.nih.gov). P.E.V. is a Fellow of MQ: Transforming Mental Health (MQF17_24). K.W. is supported by the Welcome Trust (215901/Z/19/Z). L.Z.J. is supported by the Commonwealth Scholarship Commission, United Kingdom. D.M. and A.G.T. are supported by the Intramural Research Program of the National Institute of Mental Health (NIH annual report number ZICMH002960). J.-F.M. is supported by the French Agence Nationale de la Recherche through the grants ANR-19-CE45-0022-01IFOPASUBA and ANR-20-CHIA-0027-01FOLDDICO. V.K. is supported by the MRC translation support award [MR/V036874/1] and the Developing Human Connectome Project.

We are grateful for the efforts of the teams involved in the Developing Human Connectome Project (dHCP) towards the collection and processing of the fetal neuroimaging data. The dHCP is funded by the European Research Council under the European Union’s Seventh Framework Programme (FP/2007-2013) / ERC Grant Agreement no. [319456]. We are thankful for the families that participated in this data collection as well as for the participants in the UK Biobank study for making this work possible. The UK Biobank data was obtained under application number 22875.

## Contributions

Conceptualization and Methodology, W.E.S., P.E.V., A.R., and E.T.B; Data Curation, W.E.S., V.K., K.W., L.Z.J., D.M., A.G.T., V.R.K., J.S., E.C.R.; Software implementation, W.E.S., D.R.; Formal analysis, W.E.S.; Writing – Original Draft, W.E.S., A.R., E.T.B.; Writing – Review and Editing, W.E.S., P.E.V., V.K., K.W., L.Z.J., D.M., A.G.T., V.R.K., J.S., D.R., E.C.R., J.-F.M., A.R., E.T.B.

## Declaration of interests

E.T.B. has consulted for GlaxoSmithKline, SR One, Sosei Heptares, Boehringer Ingelheim and Monument Therapeutics. E.T.B. and J.S. are co-founders and stockholders of Centile Bio Inc. All other authors declare no conflicts of interest.

