## Supplemental Figures 1-6, Supplemental Movies 1-2 for "A bipolar taxonomy of adult human brain sulcal morphology related to timing of fetal sulcation and trans-sulcal gene expression gradients"

### Contents

**Supplemental Figure 1. Adult and fetal cohort age distributions.**
**Supplemental Figure 2. Sulcal nomenclature.**
**Supplemental Figure 3. Sulcal covariation between sulci across subjects.**
**Supplemental Figure 4. Sulcal phenotypes, but not SPNs, exhibit considerable age, sex, and total brain volume effects.**
**Supplemental Figure 5. SPNs have highly reproducible estimates of sulcal similarity and clustering properties.**
**Supplemental Figure 6. The group mean SPN was best captured by a single principal component.**

**Supplemental Movie 1. Animation of linear sulci developmental emergence.**
**Supplemental Movie 2. Animation of complex sulci developmental emergence.**

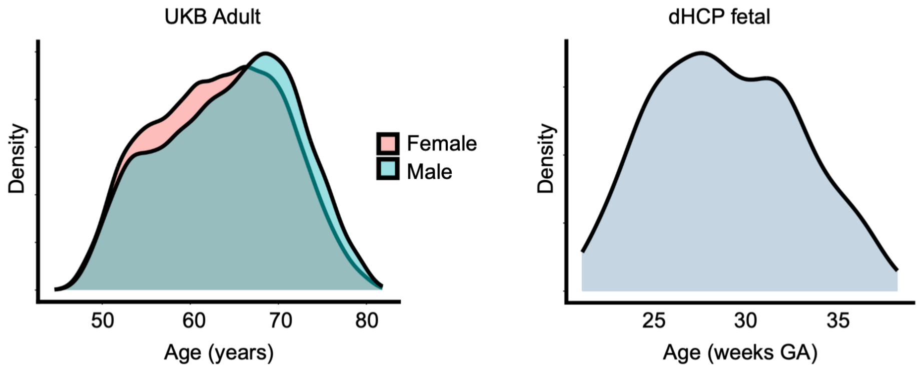


**Supplemental Figure 1. Adult and fetal cohort age distributions.** Subjects from the UK Biobank included in adult brain analyses spanned 45-82 years of age with sufficient representation of subjects to capture age related changes over this period. Subjects from the dHCP included in fetal brain analyses spanned 21-36 weeks GA. At the ends of sulcal development, where relatively lower inter-subject variability in global curvature is observed, three brains were matched to the 21 weeks GA template and four brains were matched to the 36 weeks GA template. A mean 14.25 brains were matched to each week-averaged template, enabling estimate of sulcal curvature trajectories throughout sulcal development. dHCP, developing Human Connectome Project; GA, gestational age.

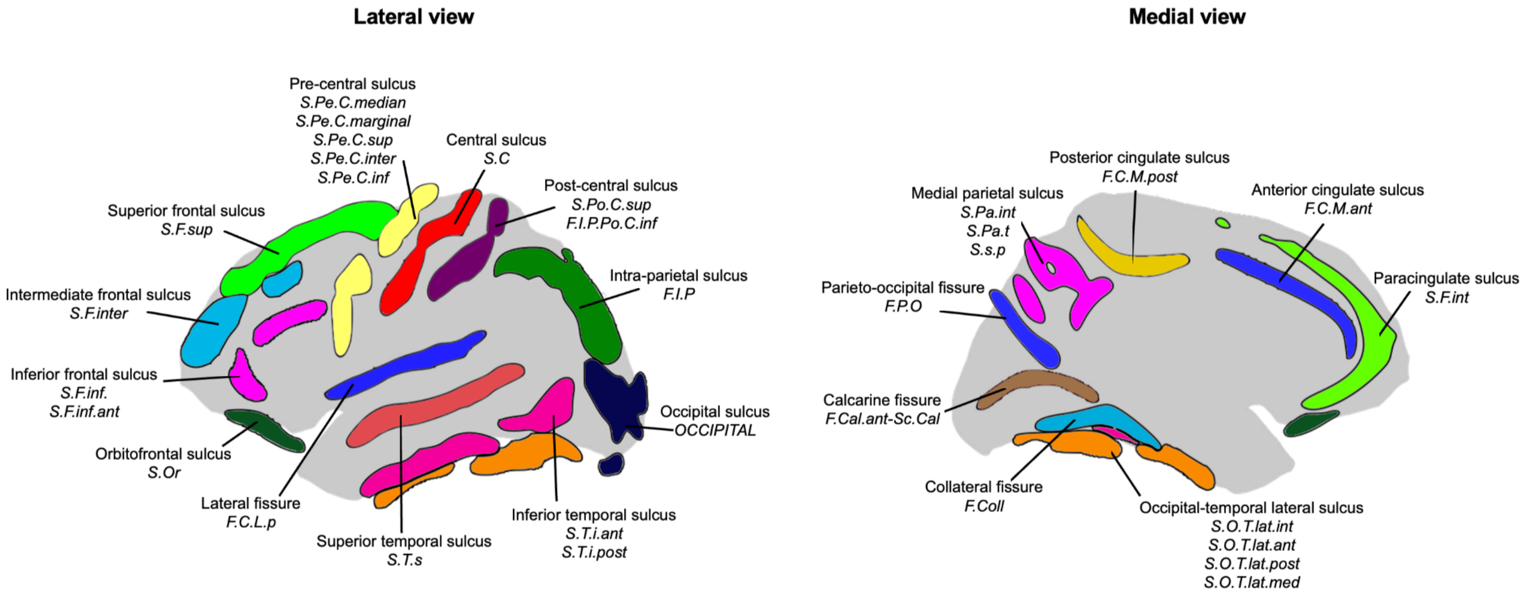


| BrainVISA label | Label in present study | BrainVISA label | Label in present study |
|---|---|---|---|
| F.Cal.ant-Sc.Cal. | Calcarine fissure | S.F.int. | **Paracingulate sulcus** |
| F.Coll. | Collateral fissure | S.F.inter. | Intermediate frontal sulcus |
| F.I.P. | Intra-parietal fissure | S.F.sup. | Superior frontal sulcus |
| F.C.L.p. | Lateral fissure | S.Or. | Orbitofrontal sulcus |
| F.C.M.ant | Anterior cingulate sulcus | S.O.T.lat.(int/ant/post/med) | Occipital-temporal lateral sulcus |
| F.C.M.post. | Posterior cingulate sulcus | S.Pa.(int/t) / S.s.p. | Medial parietal sulcus |
| OCCIPITAL | Occipital sulcus | S.Pe.C.(median/marginal/sup/inter/inf) | Pre-central sulcus |
| F.C.L.p. | Lateral fissure | S.Po.C.Sup / F.I.P.Po.C.inf | Pos-central sulcus |
| S.C. | Central sulcus | S.T.i.(ant/post) | Inferior temporal sulcus |
| S.F.inf.(ant) | Inferior frontal sulcus | S.T.s. | Superior temporal sulcus |

**Supplemental Figure 2. Sulcal nomenclature.** Lateral and medial views of the cortex are shown with sulci shown as the average path between gyri. Sulcal labels used in this study come from the standard BrainVISA Morphologist taxonomy, with the French abbreviations used by BrainVISA italicized underneath. These labeling schemes are also presented in table format.

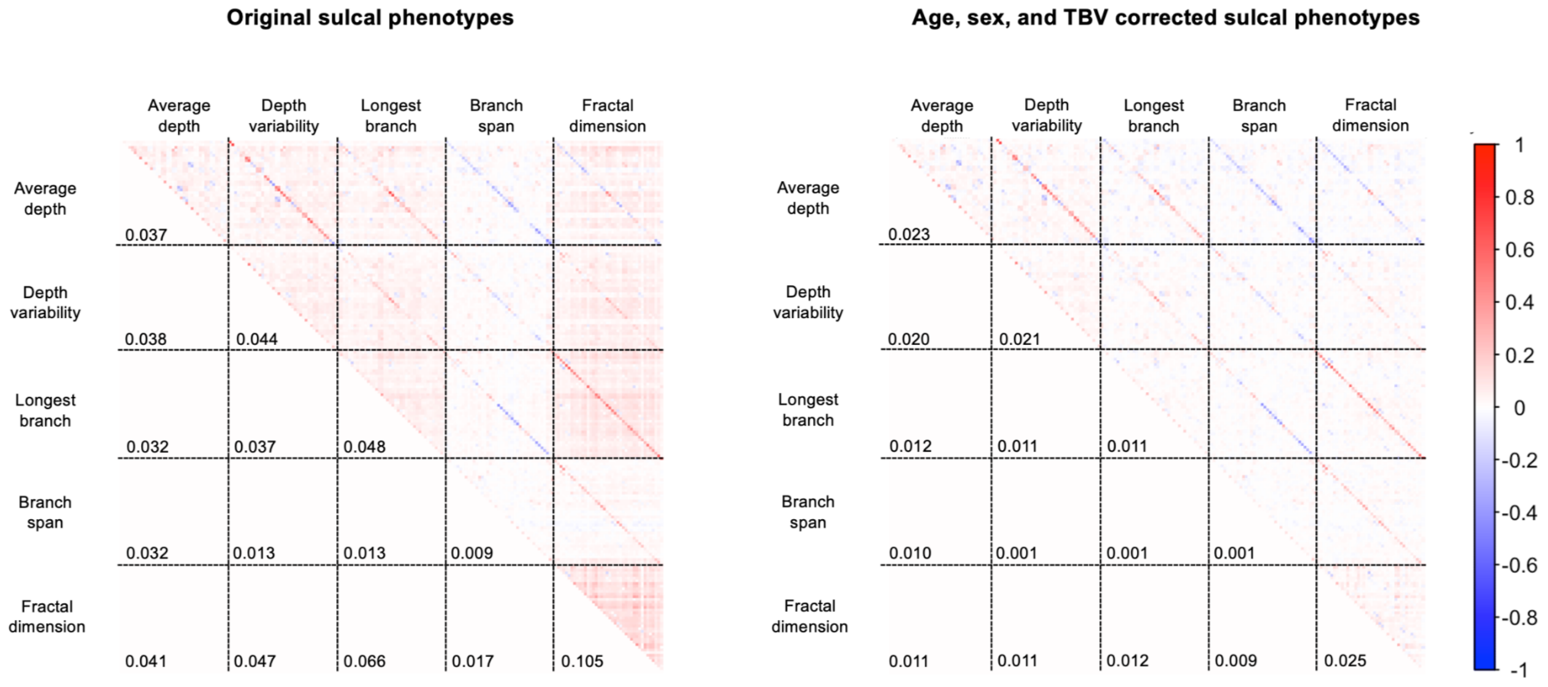


**Supplemental Figure 3. Sulcal covariation between sulci across subjects.** (Left) The correlation across subjects between any two sulci, repeated for all pairs of sulcal phenotypes, was low for any given set of sulci and sulcal phenotype (median absolute correlation = 0.03). Values on the lower triangle show the median absolute correlation between sulci across subjects for any set of two sulcal phenotypes. Diagonals within any given block represent the correlation between the same sulcus for two different phenotypes; the greater magnitude correlations along these diagonals is consistent with the idea that morphology is largely independent between sulci but forms by more consistent rules across phenotypes within a given sulcus. (Right) The analysis was repeated but for sulcal phenotype values corrected for age, sex, and TBV. Again, low correlations were observed pairwise between any two sulci for any two sulcal phenotypes (median absolute correlation = 0.01), with slightly decreased block-wise median absolute correlations as compared to uncorrected sulcal phenotypes. Of note, block-wise diagonals demonstrated low but relatively higher sulcal covariation between the same sulcus for two phenotypes (median absolute correlation = 0.17, and after residualization = 0.16). Interhemispheric correlations were also slightly above average of all correlations (median absolute correlation = 0.05, and after residualization = 0.03).

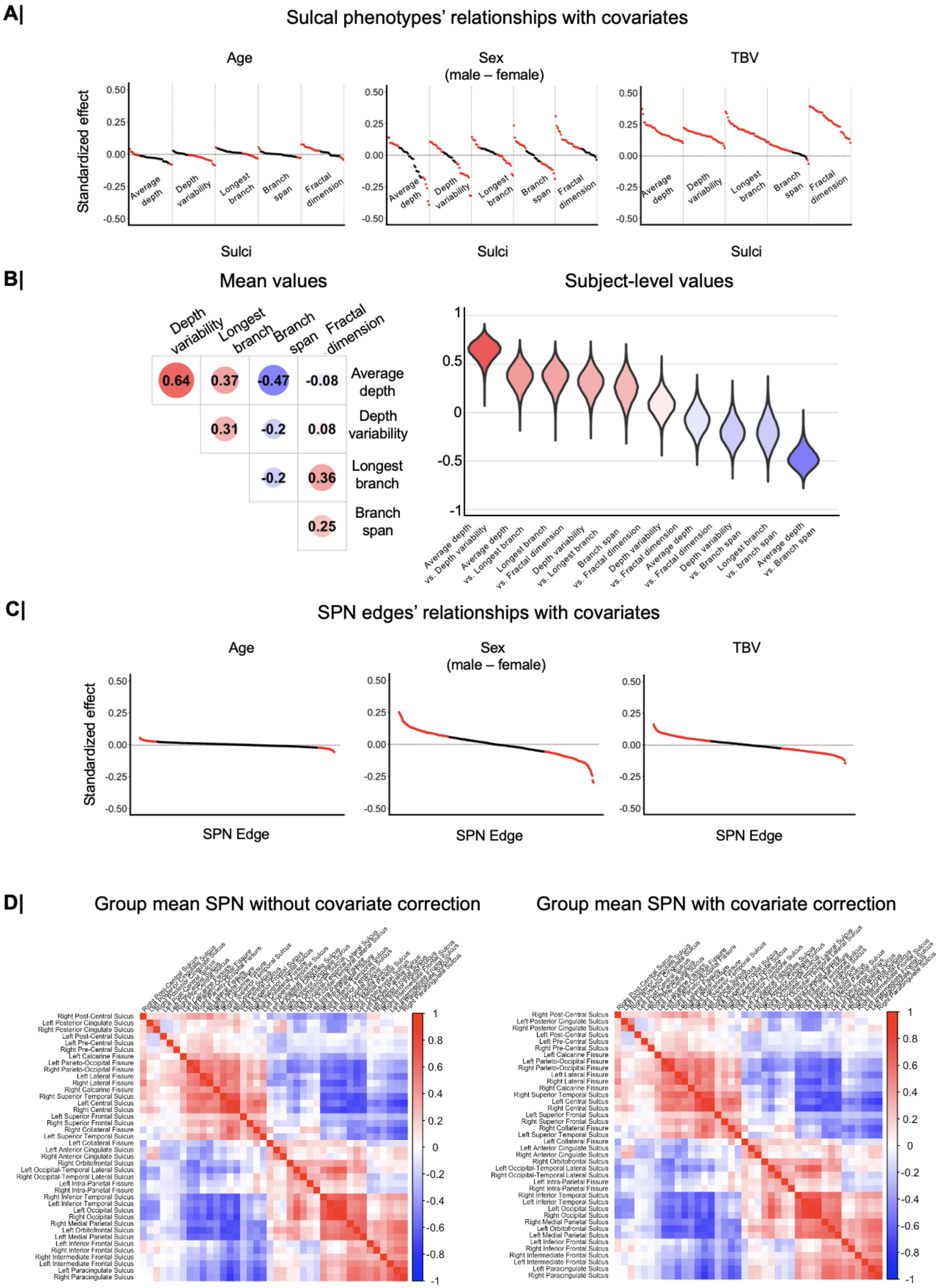
A|
Sulcal phenotypes' relationships with covariates
Age
Sex
(male – female)
TBV
Standardized effect
0.50
0.25
0.00
-0.25
-0.50
Average depth
Depth variability
Longest branch
Branch span
Fractal dimension
Sulci
B|
Mean values
Depth variability
Longest branch
Branch span
Fractal dimension
Average depth
0.64
0.37
-0.47
-0.08
0.31
-0.2
0.08
0.36
0.25
Subject-level values
1
0.5
0
-0.5
-1
Average depth vs. Depth variability
Average depth vs. Longest branch
Longest branch vs. Fractal dimension
Depth variability vs. Longest branch
Branch span vs. Fractal dimension
Depth variability vs. Fractal dimension
Average depth vs. Fractal dimension
Depth variability vs. Branch span
Longest branch vs. branch span
Average depth vs. Branch span
C|
SPN edges' relationships with covariates
SPN Edge
D|
Group mean SPN without covariate correction
Group mean SPN with covariate correction
1
0.8
0.6
0.4
0.2
0
-0.2
-0.4
-0.6
-0.8
-1

**Supplemental Figure 4. Sulcal phenotypes, but not SPNs, exhibit considerable age, sex, and total brain volume effects. (A)** First, we demonstrated age, sex, and total brain volume effects at the level of sulcal phenotypes. Dots show standardized effect size from multiple linear regression of each sulcus and phenotype with age, sex, and total brain volume. Dots were ordered by effect magnitude, with red dots indicating sulci significant after Bonferroni correction for 120 tests (40 sulci * three covariates). Age and sex displayed both positive and negative effects on any given sulcus and sulcal phenotype, whereas increased total brain volume largely corresponded to increased sulcal phenotype value for all sulci. **(B)** Analyses from **Figure 2B** were repeated with covariate residualized sulcal phenotypes given the above significant effects of age, sex, and TBV. Almost identical distributions of within-brain covariation of sulcal phenotypes were observed compared to covariation of the original, uncorrected sulcal phenotypes. **(C)** The same models as in **(A)** were instead fit to each of the 780 unique edges in the symmetric 40x40 sulcus SPN matrices. Red dots showed significance for any SPN edge (unique sulcus vs. sulcus correlation strength) after Bonferroni correction for 2340 tests (780 edges * three covariates). In general, standardized effect sizes were decreased for SPN edges as compared to prior tests with underlying sulcal phenotypes, suggesting the relative relationships of sulci are robust to age, sex, and total brain volume. **(D)** We explicitly addressed the impact of age, sex, and total brain volume on the group mean SPN that served as the basis for most downstream analyses. By regressing out the effects of age, sex, and total brain volume on sulcal phenotypes as in **(A)**, subject-level and group mean corrected SPNs were generated. Corrected subject-level SPNs had mean edge-wise correlation of 0.996 with their corresponding uncorrected SPN. We plotted the group mean SPN, corrected and uncorrected, to highlight SPN robustness to age, sex, and total brain volume given their identical structure. The uncorrected and corrected group mean SPN had edgewise correlation of approximately 1 ($r = 0.9999917$).

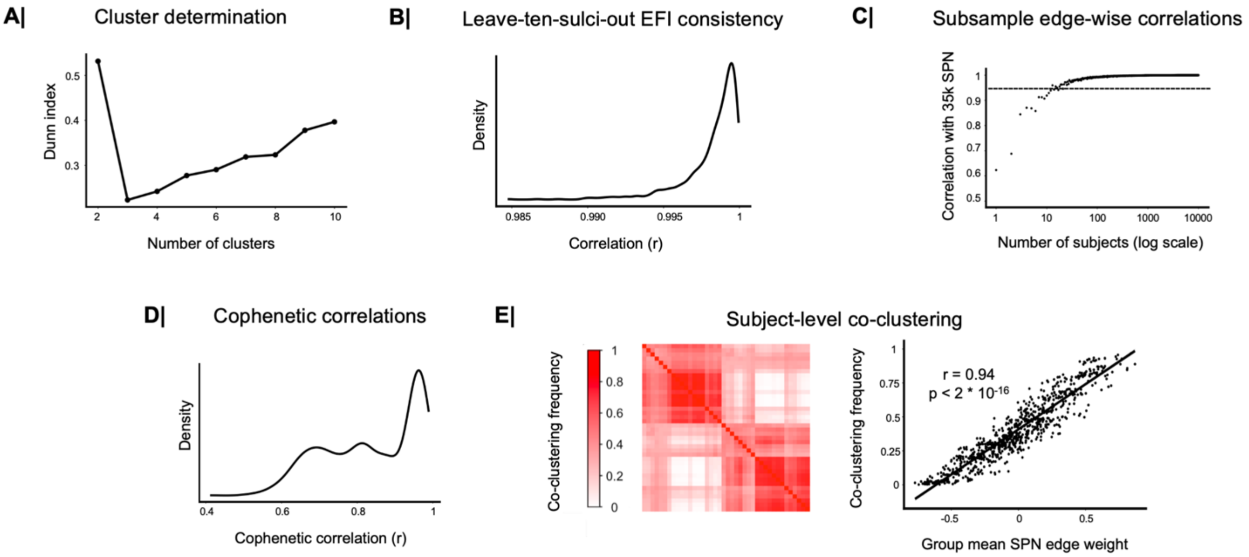


**Supplemental Figure 5. SPNs have highly reproducible estimates of sulcal similarity and clustering properties. (A)** Dunn index was calculated for the group mean SPN hierarchical clustering from clusters of size 2 through 10, with the index maximized (*i.e.*, optimal) at a two-cluster solution. (**B**) SPNs were robust to choice of sulci used in the study. The first principal component values for the group mean SPN was correlated with the first principal component for the group mean SPN with the exclusion of 10 sulci (*i.e.*, 30 remaining sulci retained) repeated for 1000 random subsets of the 10 excluded sulci. No particular subset of sulci had a large impact on the principal component structure, with r > 0.984 for all permutations. (**C**) Random samples of increasing size were pooled from all 34,725 subjects and were averaged across edges to yield subsample mean SPNs. Edge-wise correlation of subsample mean SPNs were correlated with the group (34,725 subjects) mean SPN to assess the number of subjects needed to reach our main findings from the group mean SPN. A correlation of 0.95 (dashed line) can be seen at as low as n = 15 subjects in the subsample. **(D)** We next addressed whether the hierarchical clustering of SPNs was reproducible, given our main finding of two clusters of sulcal morphology. Subsample mean SPNs were generated with random samples of n = 20 subjects to assess inter-sample variability. 1000 subsample mean SPNs were generated, clustered with hierarchical clustering, and had dendrogram distances compared to the group mean SPN (34,725 subjects) with cophenetic correlation. Clustering solutions from these small subsamples yielded high correspondence with the structure of the group mean SPN, with mean cophenetic correlation = 0.83. **(E)** Additionally, we tested whether subject-level clustering solutions had a consistent pattern. Subject SPNs were automatically clustered using the Louvain algorithm with r = 0 threshold SPNs, generating matrices giving values of 1 where two sulci were clustered together and 0 elsewhere. The average of these matrices across subjects yielded a co-clustering frequency matrix, strongly resembling the group mean SPN as assessed by edgewise correlation (r = 0.94). Therefore, we found that SPN estimates of sulcus-by-sulcus morphological similarity and related anatomical classes of linear and complex sulci were well-reflected in individuals and small samples of subjects. In additional analyses, hierarchical clustering of subject-level clustering patterns did not reveal any clusters of subjects that form separable patterns of sulcal clustering, with the overall bipartite division of SPNs remaining as the dominant clustering pattern.

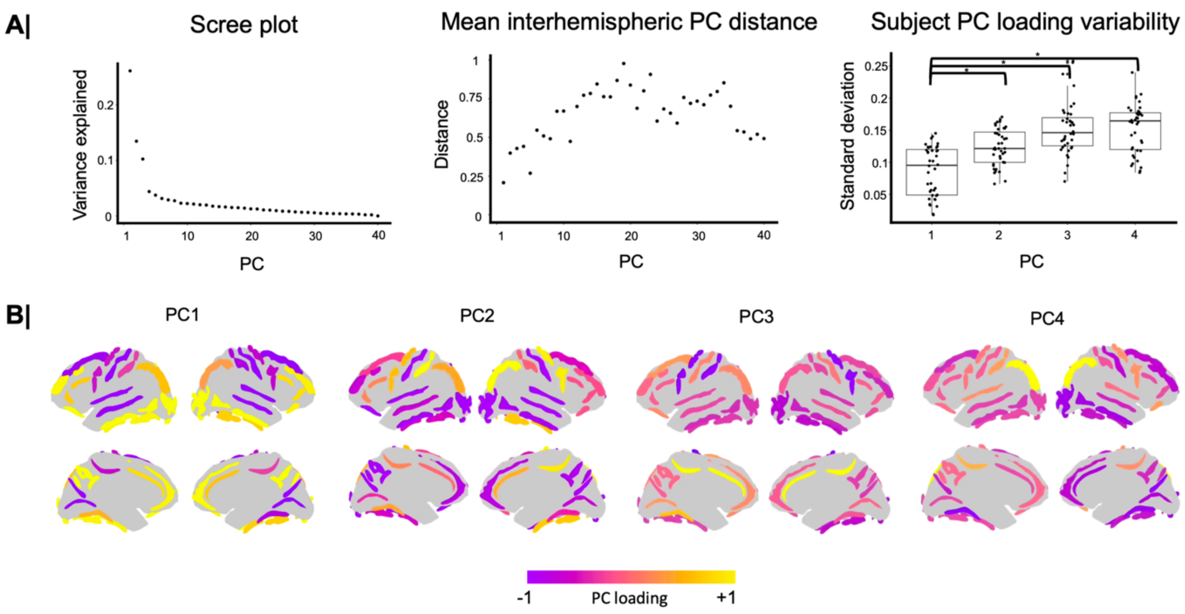


**Supplemental Figure 6. The group mean SPN was best captured by a single principal component. (A)** Evaluation of principal components of group mean SPN. A scree plot for principal component analysis of the group mean SPN showed that the first principal component dominantly explained variance (27%), with decreasing variance explained by further components. To assess whether components explaining less variance were biologically meaningful, the interhemispheric distance along standardized principal component (PC) loadings for each PC were assessed. Large interhemispheric differences could indicate a noisy latent dimension that is not reproducible across hemispheres. Interhemispheric PC distance increased with PC number, with comparable distance to PC1 only achieved at a PC that explained less than 5% variance. Lateralized PC loadings could still be biologically meaningful, however, so we tested additionally for consistency of PC loadings across subjects, plotting the standard deviation for each sulcus' PC loading across subjects from subject level principal component decomposition. The first four PCs plotted, demonstrating PC loading variability for PCs up to the elbow in the scree plot. PCs 2, 3, and 4 all had significantly greater ($p < 0.001$) standard deviation in sulcus PC loadings than PC1. **(B)** We additionally plotted each of the top four PCs for visual inspection. The greater interhemispheric similarity for PC1 relative to other PCs is readily visible. Additional PCs had loadings that were not well differentiated amongst sulci except for few outliers.

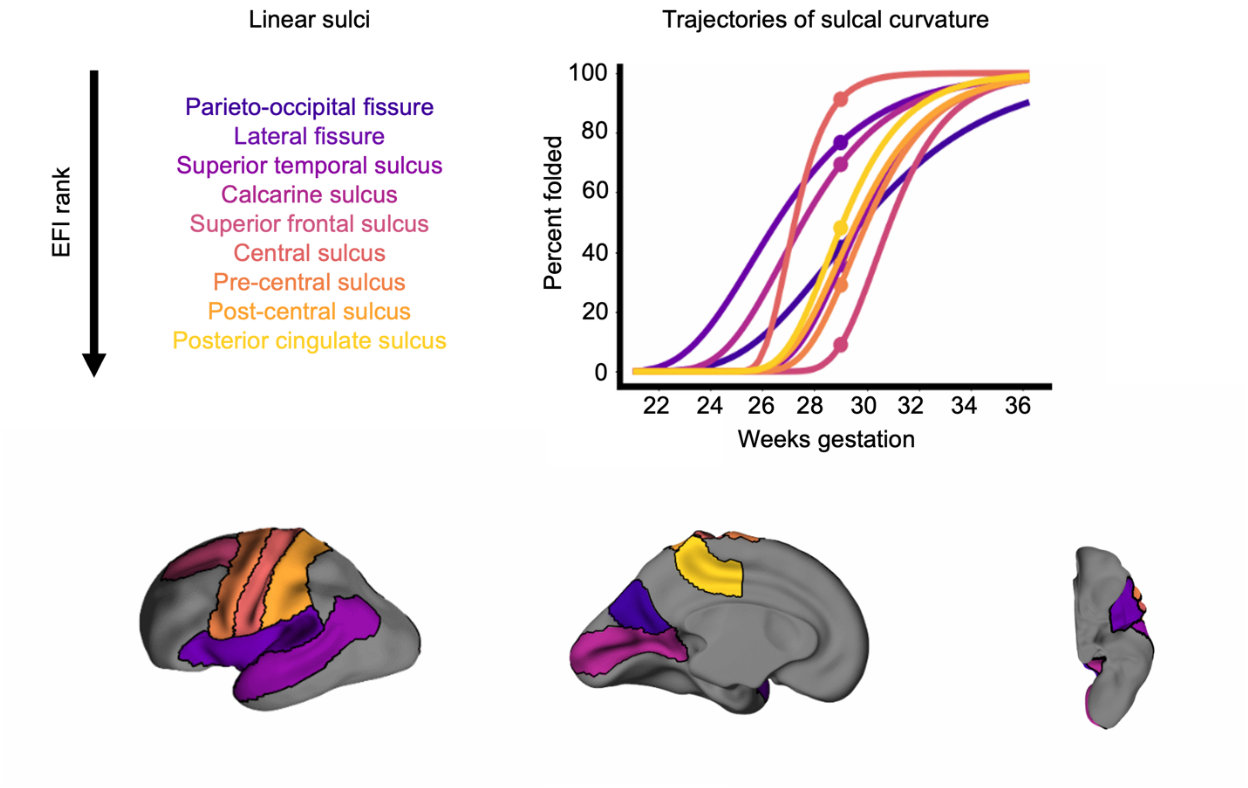


https://youtu.be/fmQ3TEMDU1o

**Supplemental Movie 1. Animation of linear sulci developmental emergence.** (Use the link to play video on YouTube). Linear sulci were colored by their relative EFI rank within the linear cluster to distinguish the sulcal trajectories and the sulcal regions mapped with the same colors. Lateral, medial, and ventral views of the cortex were temporally interpolated between gestation-week templates of the fetal cortical surface between 21 and 36 gestational weeks. The movie shows the agreement between the models of sulcal development (top right) and the visual appearance of in-folding cortical sulci (bottom). For example, the calcarine and parieto-occipital fissure (respectively, the second and third sulci to emerge at 10% folded), can both be seen to indent early relative to other regions, yet this indentation occurs much more rapidly in the calcarine fissure. This is reflected in the greater rate of folding in the model fit for the calcarine sulcus. Additionally, it is useful to focus on all color-labeled and all non-labeled sulci to see how folds form first in the labeled regions.

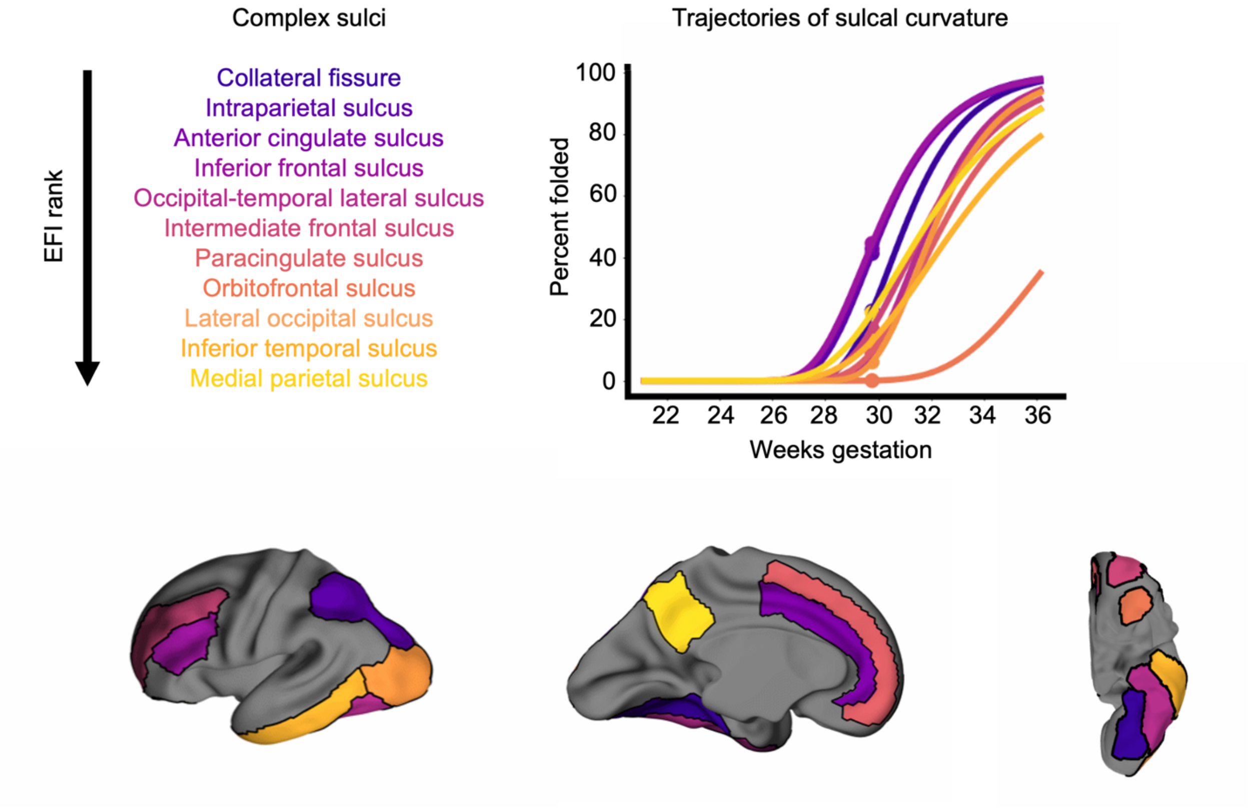


https://youtu.be/IcGRXZTy5Ic

**Supplemental Movie 2. Animation of complex sulci developmental emergence.** (Use the link to play video on YouTube). Complex sulci are colored by their relative EFI rank within the complex cluster to distinguish the sulcal trajectories and the sulcal regions mapped with the same colors. Lateral, medial, and ventral views of the cortex again were animated over the 21-36 gestational week period. The models fit to complex sulci are also validated in the animation, such as seen with the orbitofrontal sulcus (the last sulcus to form). For this sulcus, its ventral positioning shifts as more tissue distinguishes the frontal versus ventral regions of cortex, but indentation does not occur until late in development (~33 weeks gestation). Again, this observation is reflected in the model fit for the orbitofrontal sulcus. By observing all color-labeled regions versus all non-labeled regions, it can be seen that the non-labeled regions fill in the cortical surface with folds prior to the emergence of complex sulci.
